# Circadian clock disruption and growth of kidney cysts in autosomal dominant polycystic kidney disease

**DOI:** 10.1101/2024.08.05.606676

**Authors:** Abeda Jamadar, Christopher J. Ward, Viji Remadevi, Meekha M Varghese, Navjot S Pabla, Michelle L. Gumz, Reena Rao

**Affiliations:** Kidney Institute, University of Kansas Medical Center, Kansas City, KS. USA; Department of Medicine, Division of Nephrology, University of Kansas Medical Center, Kansas City, KS. USA; Division of Pharmaceutics and Pharmacology, College of Pharmacy & Comprehensive Cancer Center, The Ohio State University, Columbus, Ohio, USA; Department of Physiology and Aging, Department of Medicine, Division of Nephrology, Hypertension, and Renal Transplantation, University of Florida, Gainesville, FL, USA

**Keywords:** Circadian clock, BMAL1, cell proliferation, cyst, lipogenesis, fatty-acid oxidation

## Abstract

**Background:** Autosomal dominant polycystic kidney disease (ADPKD) is caused by mutations in the *PKD1* and *PKD2* genes, and often progresses to kidney failure. ADPKD progression is not uniform among patients, suggesting that factors secondary to the *PKD1/2* gene mutation could regulate the rate of disease progression. Here we tested the effect of circadian clock disruption on ADPKD progression. Circadian rhythms are regulated by cell-autonomous circadian clocks composed of clock proteins. BMAL1 is a core constituent of the circadian clock.

**Methods:** To disrupt the circadian clock, we deleted *Bmal1* gene in the renal collecting ducts of the *Pkd1*^RC/RC^ (RC/RC) mouse model of ADPKD (RC/RC;*Bmal1*^f/f^;*Pkhd1*^cre^, called DKO mice), and in *Pkd1* knockout mouse inner medullary collecting duct cells (*Pkd1Bmal1*KO mIMCD3 cells). Only male mice were used.

**Results:** Human nephrectomy ADPKD kidneys and *Pkd1*KO mIMCD3 cells showed reduced *Bmal1* gene expression compared to normal controls. When compared to RC/RC kidneys, DKO kidneys showed significantly altered clock gene expression, increased cyst growth, cell proliferation, apoptosis and fibrosis. DKO kidneys also showed increased lipogenesis and cholesterol synthesis-related gene expression, and increased tissue triglyceride levels compared to RC/RC kidneys. Similarly, *in vitro, Pkd1Bmal1*KO cells showed altered clock genes, increased lipogenesis and cholesterol synthesis-related genes, and reduced fatty-acid oxidation-related gene expression compared to *Pkd1KO* cells. The *Pkd1Bmal1*KO cells showed increased cell proliferation compared to *Pkd1KO* cells, which was rescued by pharmacological inhibition of lipogenesis.

**Conclusion:** Renal collecting duct specific *Bmal1* gene deletion disrupts the circadian clock and triggers accelerated ADPKD progression by altering lipid metabolism-related gene expression.

**Key points:**

- Lack of BMAL1, a circadian clock protein in renal collecting ducts disrupted the clock and increased cyst growth and fibrosis in an ADPKD mouse model.
- BMAL1 gene deletion increased cell proliferation by increasing lipogenesis in kidney cells.
- Thus, circadian clock disruption could be a risk factor for accelerated disease progression in patients with ADPKD.

## Introduction

Autosomal dominant polycystic kidney disease (ADPKD) is the most common inherited kidney disease which is caused by mutations in the *PKD1* and *PKD2* genes. Progressive cyst growth in ADPKD kidneys is accompanied by inflammation and fibrosis, which often leads to loss of renal function and kidney failure. However, disease progression to kidney failure varies significantly among ADPKD patients, even within families with the same *PKD1* gene mutation ^1^. Based on the Mayo-Irazabal classification for age-adjusted assessment of ADPKD progression, the predicted incidence of kidney failure at 10 years ranges widely between 2.2 to 77.4% in older adults ^2^. Thus, factors secondary to the *PKD1/2* gene mutation could regulate the rate of ADPKD progression to kidney failure. Here we tested if disruption of the circadian clock is a trigger for accelerated ADPKD progression.

Circadian rhythms are intrinsic, cyclical ∼24-hour oscillations in behavior and physiology that coordinate biological processes with the time of day. In mammals, circadian rhythms are regulated by cell-autonomous circadian clocks, which at the molecular level are comprised of clock proteins. The central clock, located in the suprachiasmatic nucleus in the hypothalamus, synchronizes peripheral clocks based on external timing cues. The canonical circadian clock involves heterodimerization of circadian proteins (transcription factors), CLOCK and BMAL1 (activators), which activate the gene expression of *Per1,2,3* and *Cry1,2* (repressors). PER and CRY proteins accumulate over time, repressing CLOCK and BMAL1 action, thereby inhibiting their own transcription. RORα/β and REV-ERBα/β function in an ancillary loop to mediate opposing action on BMAL1 transcription, thus driving rhythmic expression of BMAL1. These feedback loops result in ∼24-hour oscillations of clock gene expression, and rhythmic expression of ∼40% of protein-coding genes in mammals ^3^.

In mouse kidneys, ∼13% of all genes are expressed in a circadian manner ^4, 5^. Moreover, kidney functions including blood pressure rhythms, renal blood flow, glomerular filtration rate (GFR), and excretion of electrolytes such as sodium and potassium display circadian influence ^6-9^. Chronic disruption of the circadian clock (chronodisruption) is a well-known risk factor for accelerated ageing, memory loss, metabolic defects, obesity, cancer, cardiovascular defects, and metabolic syndrome in humans ^10, 11^. The role of chronodisruption in ADPKD pathogenesis is currently unclear.

BMAL1 is encoded by the *Arntl* (Aryl hydrocarbon receptor nuclear translocator-like protein-1) gene and is a core component of the molecular circadian clock. Together with other clock proteins, BMAL1 plays an essential role in cellular metabolic processes by regulating time-dependent utilization of nutrients and metabolites to support cell growth ^12^. *Bmal1* is a master clock gene, and the only clock gene whose deletion alone leads to complete loss of circadian rhythms ^13^. Here we examined the effect of collecting duct-specific gene deletion of *Bmal1* on disease progression in an ADPKD mouse model and in *Pkd1* gene knockout cells.

## Methods

### Human tissue samples

De-identified human normal and APKD kidney tissues were “medical discard” nephrectomy samples from the Kansas PKD Research and Translational Center. IRB approval #5929, 8/25/2020. The time of tissue collection is unknown.

### *Bmal1* gene knockout in WT and *Pkd1*KO mIMCD3 cells using Crispr-Cas12a

Murine *Bmal1* (*Arntl* gene, ID 11865) was disrupted in mIMCD3 cells using a microhomology end joining-based strategy and Cas12a ^14^. The **TTTG**GCGTATCTACCACAGGAACT sequence in exon 13 was targeted (PAM in bold). CRISPR sequence was, 5’/AltR1/rUrArArUrUrUrCrUrArCrUrCrUrUrGrUrArGrArUrGrCrGrUrArUrCrUrArCrCrArCrArGrGrA rArCrU/AltR2/-3’. PCR products that showed deletion on PAGE were sequenced after alkaline phosphatase treatment (ExoSAP, # 78200, Applied Biosystems, Foster City, CA). *Bmal1*KO clone and *Pkd1Bmal1*KO had frame-shift deletions of 10bp and 16bp respectively.

### Mouse model

RC/RC (p.R3277C) mice ^15^ were bred with *Bmal1*^f/f^ mice (B6.129S4(Cg)- *Bmal1tm1Weit/J*, carrying loxP sites flanking exon 8 of *Bmal1* gene, Strain#007668, The Jackson Laboratory), and mice carrying Pkhd1^cre 16^. The resultant *Pkd1*^RC/RC^;*Bmal1*^f/f^;*Pkhd1*^cre^ mice (double knockout DKO) were compared with *Bmal1*^f/f^ (wild type-WT), *Bmal1*^f/f^;*Pkhd1*^cre^ (*Bmal1*KO) and RC/RC littermates. Only male mice were used and sacrificed at 8 months of age, between 11:30 AM and 12:30 PM. Studies were approved by our Univerisity’s IACUC committee.

### Recombination specificity in *Bmal1*KO and DKO mice

*Bmal1* recombination was confirmed by PCR on genomic DNA of whole papilla from mouse kidneys ^17, 18^. PCR reaction products run on a 2% agarose gel yielded a 431-bp band for floxed *Bmal1* allele and 572 bp band for recombination product.

### Statistics

Values are expressed as mean±SEM for *in vivo* and mean±SD for *in vitro* studies. Data was analyzed by two-tailed unpaired T-test with Welch’s correction; or one-way ANOVA followed by Tukey’s multiple comparison test and two-way ANOVA followed by Bonferroni’s multiple comparison test using GraphPad Prism software (Version 9). P≤0.05 was considered significant.

### Supplementary methods

We provide detailed methods for *in vitro* BrdU incorporation ^19, 20^, MTT assay ^21^, BUN ^22^ quantification of cyst ^22^, Western blot ^23, 24^, TUNEL staining ^21, 25^, immunohistochemistry/immunofluorescence (IHC/IF) staining ^26^ and quantitative real time PCR (QRTPCR) ^27^ in supplementary methods, and primer list for QRTPCR in Supplemental Table 1.

## Results

### 1. Renal collecting duct specific *Bmal1* gene deletion in ADPKD mouse kidneys

Examination of *BMAL1* (*ARNTL* gene) expression in human nephrectomy ADPKD kidneys showed over 2-fold decrease in *BMAL1* mRNA levels, compared to normal human kidneys (Fig 1A). Consistent with this, the kidney interactive transcriptomics database ^28^ of single cell RNA-sequencing showed reduced *Bmal1* expression in most renal cells of human ADPKD kidneys compared to normal control kidneys (Fig 1B). To examine the effect of *Pkd1* gene deletion on *Bmal1*, we examined its mRNA levels in control and *Pkd1*^-/-^ mouse renal inner medullary collecting duct cells (mIMCD3) ^29^ and mouse proximal tubular cells ^16, 30^. Synchronized cells were collected every 4h, for 48h. The control *Pkd1*^+/+^ mIMCD3 cells and *Pkd1*^+/-^ proximal tubule cells showed clear circadian oscillation of *Bmal1* mRNA, with ∼20h interval between peaks, while *Bmal1* mRNA levels and circadian oscillation were significantly dampened in the corresponding *Pkd1*^-/-^ cells (Fig 1C, D).

**Figure 1.**
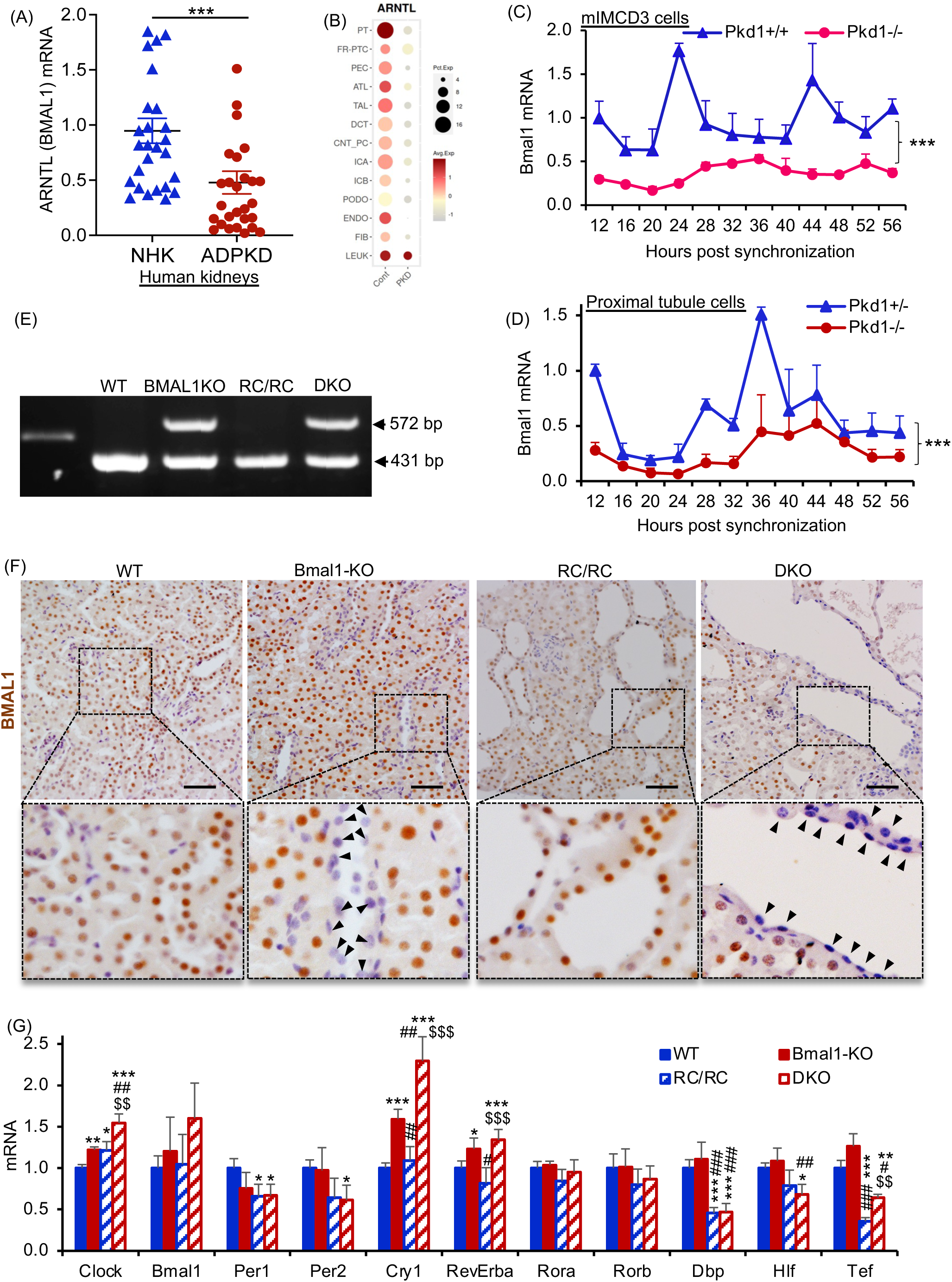
Renal collecting duct specific *Bmal1* gene knockout in WT and RC/RC mice. (A) *Bmal1* mRNA relative to 18S mRNA in normal human kidneys (NHK) and ADPKD kidneys measured by QRTPCR. ***P<0.001 by T-test. (B) *Bmal1* gene expression in human normal control and ADPKD kidneys using kidney interactive transcriptomics database of single cell RNA-seq data. (C) Time course showing fold change in *Bmal1* mRNA relative to 18S mRNA in mouse inner medullary collecting duct cells (mIMCD3) (n=4), and (D) proximal tubule cells (n=3). ***P<0.001 by 2-way ANOVA for C and D. Cells were synchronized using dexamethasone (100nM, for 2h) and released into media with 5% FBS. After 12h, cell lysates were collected every 4 hours for 48h. (E) Agarose gel showing recombination PCR products of genomic DNA obtained from mouse kidney. The 431 bp band denotes the floxed *Bmal1* allele and the 572 bp band represents the recombination product. A single 431 bp band indicates absence of recombination. (F) Immunostaining for BMAL1 (brown) in kidney tissue sections of WT, *Bmal1*KO, RC/RC and RC/RC;*Bmal1*KO (DKO) mice (male, 8 months old) (Scale bar = 100μm). (G) QRTPCR analysis of mRNA levels of clock genes relative to 18S mRNA in whole kidney tissue lysates. *=*vs* WT, # =*vs Bmal1*KO and $=*vs* RC/RC. *P<0.05, **P<0.01, ***P<0.001 by One-way ANOVA. n=5 in WT and *Bmal1*KO, and n=7 in RC/RC and DKO.

To determine the role of BMAL1 in ADPKD progression, we chose the *Pkd1* gene hypomorphic *Pkd1*^RC/RC^ (RC/RC) mouse model of ADPKD. The RC/RC mice showed no significant difference in renal *Bmal1* levels at 4 or 8 months of age compared to wild type (WT) littermates (Supplemental Fig 1A,B,C,D,E). It is possible that total loss of Pkd1 gene expression is required for Bmal1 expression to decrease. Renal collecting duct specific *Bmal1* gene knockout in WT mice- *Bmal1*^f/f^;*Pkhd1*^cre^ (*Bmal1*KO) and RC/RC mice-RC/RC;*Bmal1*^f/f^;*Pkhd1*^cre^ (DKO) mice were generated. To test the effect of *Bmal1* gene deletion, the DKO mice were compared with WT (*Bmal1*^f/f^), RC/RC and *Bmal1*KO littermates. Only male mice were used. We detected recombination of exon-8 of the *Bmal1* gene (disrupted allele) in the *Bmal1*KO and DKO kidneys, while recombination was not detected in the WT or RC/RC kidneys (Figure 1E). BMAL1 expression was detected in the nuclei of tubular epithelial cells in WT and RC/RC mouse kidneys (Fig 1F). In the *Bmal1*KO and DKO kidneys, BMAL1 was not detected in some tubular or cyst epithelial cells (Fig 1F). Most large cysts in DKO kidneys showed Dolichos biflorus agglutinin (DBA) staining, implying collecting ducts (Supplemental Fig 1F).

To determine the effect of Bmal1 gene deletion on the circadian clock, we measured clock gene expression. When compared to RC/RC kidneys, the DKO kidneys showed significantly increased mRNA levels of *Clock*, *Cry1*, *RevErba* and *Tef* (Fig 1G). RC/RC kidneys showed significant decreases in *Per1*, *Dbp* and *Tef*, and increases in *Clock* mRNA levels when compared to WT kidneys (Fig 1G). *Clock*, *Cry1* and *RevErba* genes were upregulated in both *Bmal1*KO and DKO kidneys compared to WT kidneys and RC/RC kidneys respectively (Fig 1G).

### 2. Renal collecting duct specific *Bmal1* gene deletion increased cyst growth in ADPKD mouse kidneys

At 8 months of age, the DKO kidneys were larger and more cystic (Fig 2A, 2B), with significantly higher cyst number (Fig 2C), cystic index (Fig 2D), kidney weight (Fig 2E) and kidney to body weight ratio (Fig 2F) compared to RC/RC kidneys. No significant changes were observed in the *Bmal1*KO kidneys compared to WT kidneys (Fig 2A,E,F). Morever, no change was observed in the BUN (blood urea nitrogen) levels in DKO mice when compared to RC/RC, WT and BMAL1KO mice (Supplementary Fig 1G). The results show that gene deletion of *Bmal1* in collecting ducts accelerates renal cyst growth in the ADPKD male mouse kidney but has no overt effect on WT kidneys.

**Figure 2.**
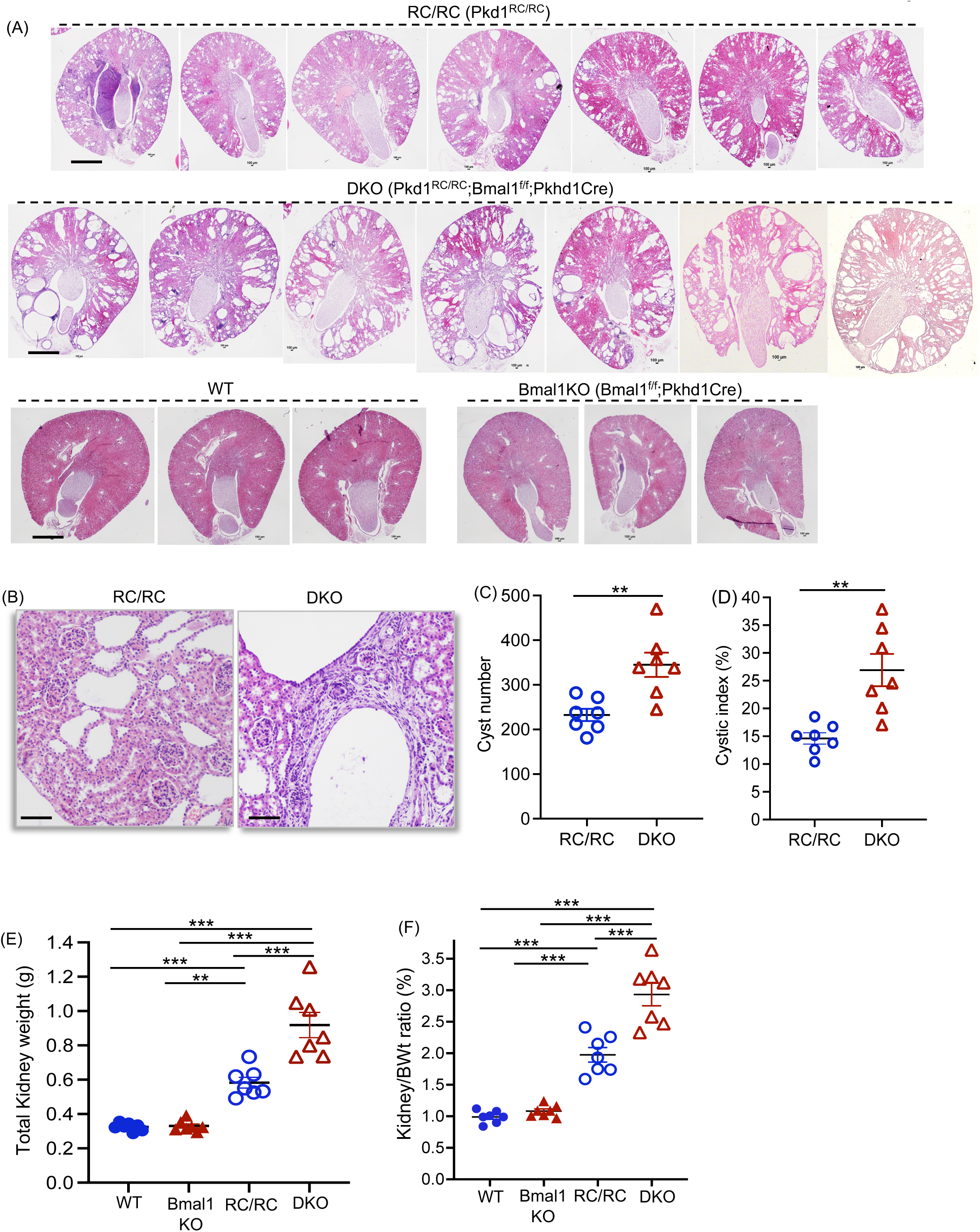
Renal collecting duct specific *Bmal1* gene knockout increased cyst growth in RC/RC mice. (A) Images of H&E staining of kidney sections and (B) magnified images. Scale bar = 1 mm for A and 400µM for B. (C) Cyst number, (D) cystic index, (E) total kidney weight, and (F) two kidney to body-weight ratio. **P<0.01, ***P<0.001 by T-test for C and D, and One-way ANOVA for E and F.

### 3. *Bmal1* gene deletion increased cell turnover and fibrosis in ADPKD mouse kidneys

To examine the effect of *Bmal1* gene deletion on cell proliferation, kidney tissue sections were immunostained for Ki67 and quantified. DKO mouse kidneys showed significantly increased Ki67 stained nuclei compared to RC/RC kidneys (Fig 3A,B). Immunoblot of kidney tissue lysate showed increased cyclin D1 and reduced P53 protein levels in DKO kidneys compared to RC/RC kidneys (Fig 3C,D,E). In DKO kidneys, pERK/ERK, pCREB/CREB, and pS6/S6 ratios were significantly increased, suggesting higher activity compared to RC/RC kidneys (Fig 3C,F,G,H). No change was observed in c-Myc levels (Supplemental Fig 2A). No significant difference was observed between *Bmal1*KO and WT kidneys in Ki67 staining (Supplemental Fig 2B, C), or in cyclin D1, pERK/ERK or pCREB/CREB protein levels (Supplemental Fig 2D,E). TUNEL staining of kidney tissue sections (Fig 4A,B) and mRNA levels of renal injury markers, *Kim1* and *Ngal1* (Fig 4C,D) were significantly increased in DKO kidneys compared to RC/RC kidneys. The increased cell proliferation and apoptosis indicate high turnover of cells in the DKO kidneys.

**Fig 3.**
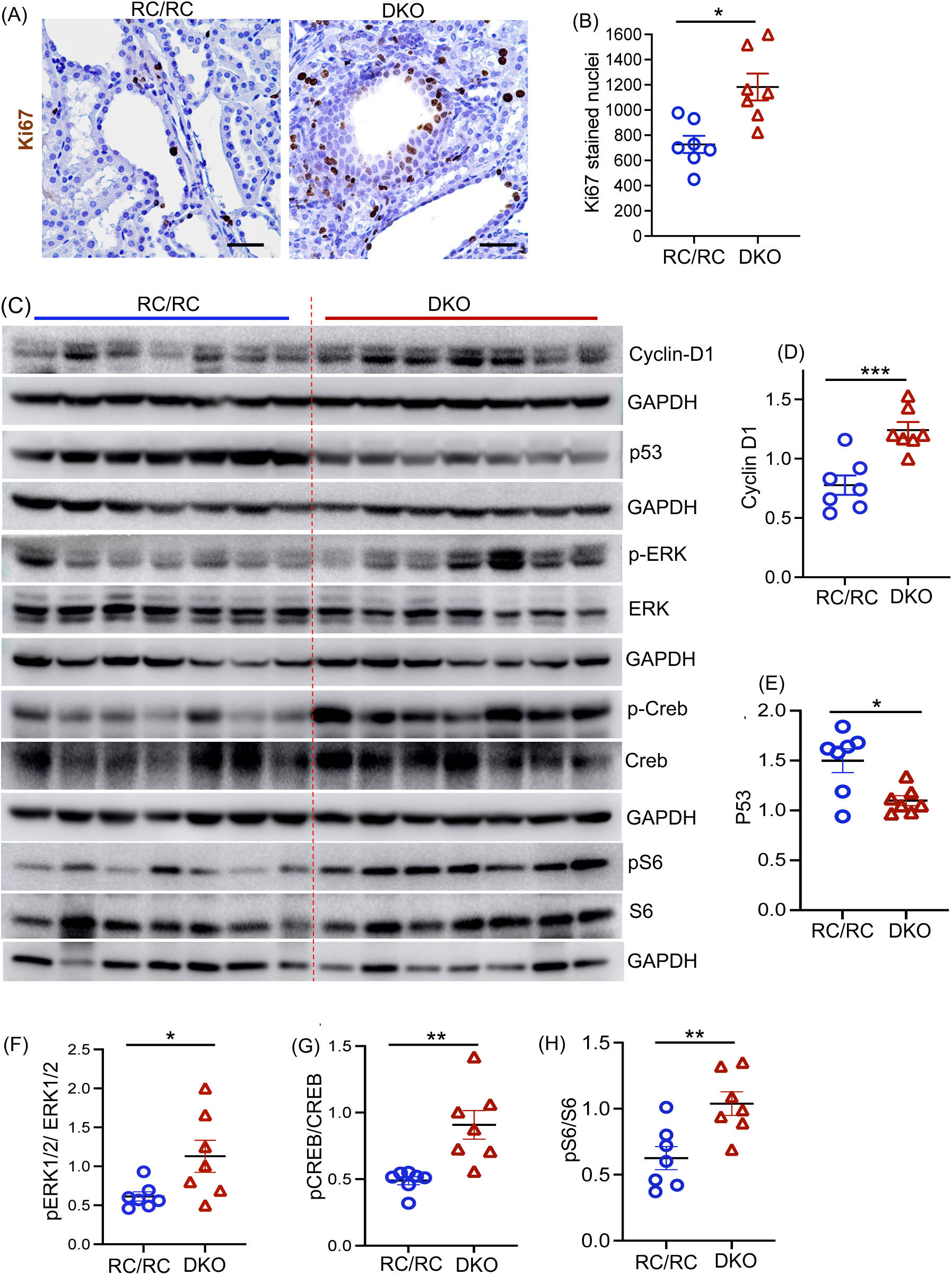
Renal collecting duct specific *Bmal1* gene knockout increased cell proliferation in RC/RC mouse kidneys. (A) Immunostaining for Ki-67 in mouse kidney tissue sections (brown nuclear staining denotes Ki-67). Scale bar = 100 µm. (B) Quantitation of Ki-67-stained nuclei per kidney section. (C) Immunoblot on kidney tissue lysate and quantitation of band density for (D) Cyclin D1, (E) P53, (F) pERK/ERK ratio, (G) pCREB/CREB ratio, (H) pS6/S6 ratio. *P<0.05, **P<0.01, ***P<0.001 by T-test.

**Fig 4.**
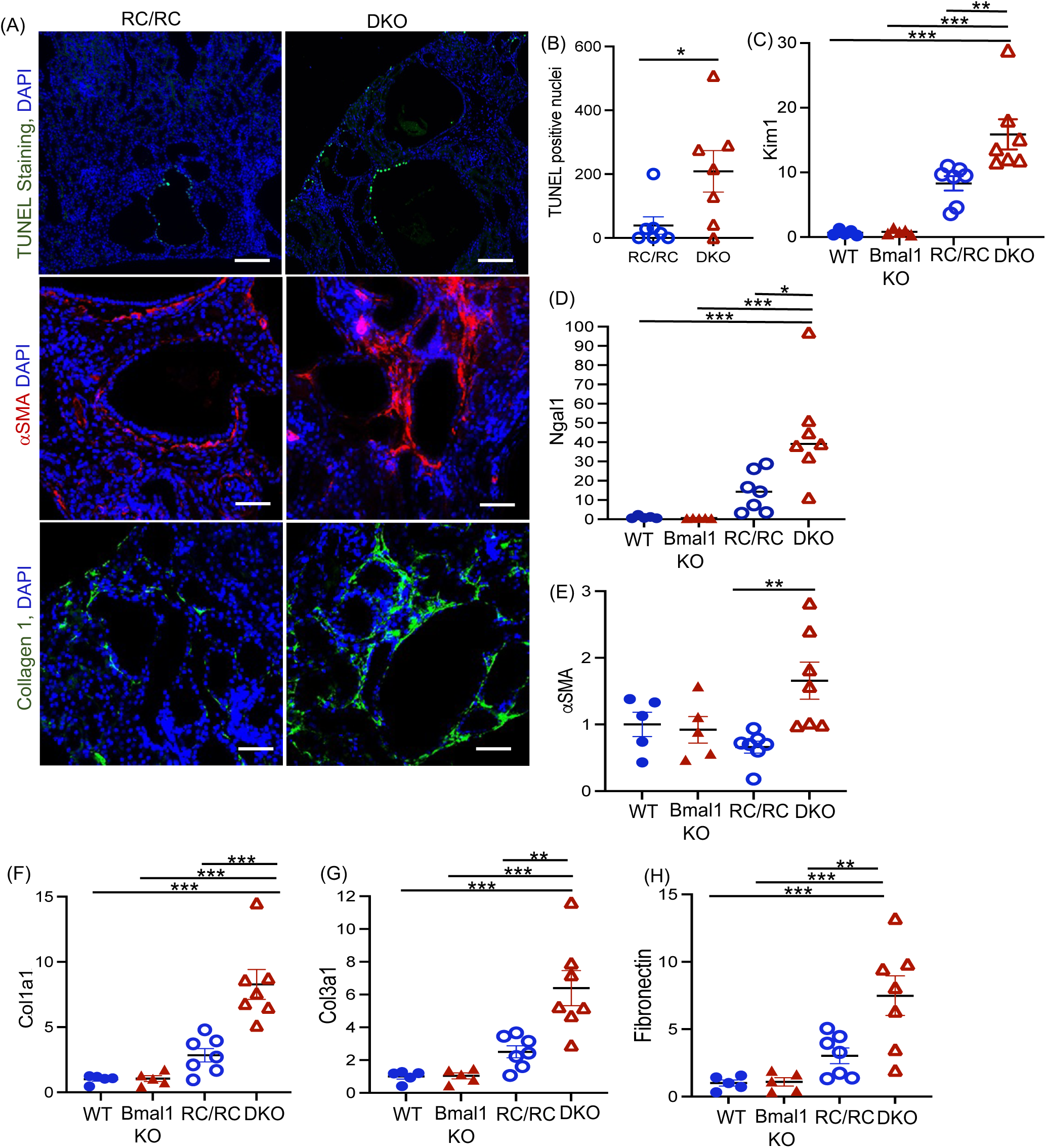
Renal collecting duct specific *Bmal1* gene knockout increased apoptosis and fibrosis in RC/RC mouse kidneys. (A) TUNEL staining (green) (Scale bar 400µm), and immunostaining for αSMA (red), and Collagen1 (green) (Scale bar 100µm) in mouse kidney tissue sections. (B) Quantitation of TUNEL staining per kidney tissue section. (C) QRTPCR for *Kim1*, (D) *Ngal1* (E) *αSMA*, (F) *Collagen 1a1*, (G) *Collagen 3a1,* and (H) *Fibronectin* in kidney tissue lysates. *P<0.05, **P<0.01, ***P<0.001 by T-test in B and One-way ANOVA in C, D, E, F, G and H.

DKO mouse kidneys also showed increased renal fibrosis. Immunostaining for the myofibroblast marker αSMA and the extracellular matrix (ECM) protein, collagen1 were increased in the DKO kidneys compared to RC/RC kidneys (Fig 4A). Moreover, mRNA levels of ECM proteins, collagen1a, collagen3a1 and fibronectin were significantly higher in the DKO kidneys compared to RC/RC kidneys (Fig 4E, F, G, H).

### 4. *Bmal1* gene knockout altered the expression of circadian clock genes and gene targets of BMAL1 in mIMCD3 cells

To better understand the mechanism by which *Bmal1* gene knockout in the renal collecting ducts accelerates disease progression in RC/RC male mice, we deleted *Bmal1* gene in WT and *Pkd1* gene knockout mIMCD3 cell lines using Crispr-Cas12a (Fig 5A, Supplemental Fig 3). WT, *Bmal1*KO, *Pkd1*KO, and *Pkd1Bmal1*KO cells were compared. Consistent with results in mouse kidneys, cell proliferation was significantly increased in *Pkd1Bmal1*KO cells compared to *Pkd1*KO cells, while *Bmal1*KO cells showed no significant difference compared to WT cells (Fig 5B). Examination of circadian clock gene expression showed significantly reduced *Per1*, *Per2*, *RevErba* and *Rora*, and increased *Cry1* mRNA levels in *Pkd1Bmal1*KO cells when compared to *Pkd1*KO cells (Fig 5C). The *Pkd1*KO cells showed significant increase in *Per1, Per2*, *RevErba, Rora* and *Hlf* mRNA levels compared to WT cells (Fig 5C). Significant difference in *Per2*, *Cry1* and *RevErba*, members of the repressor arm of the clock were common to both *Bmal1*KO and *Pkd1Bmal1*KO cells when compared to WT and *Pkd1*KO cells respectively (Fig 5C). *Rorb* and *Dbp* were not detected.

**Fig 5.**
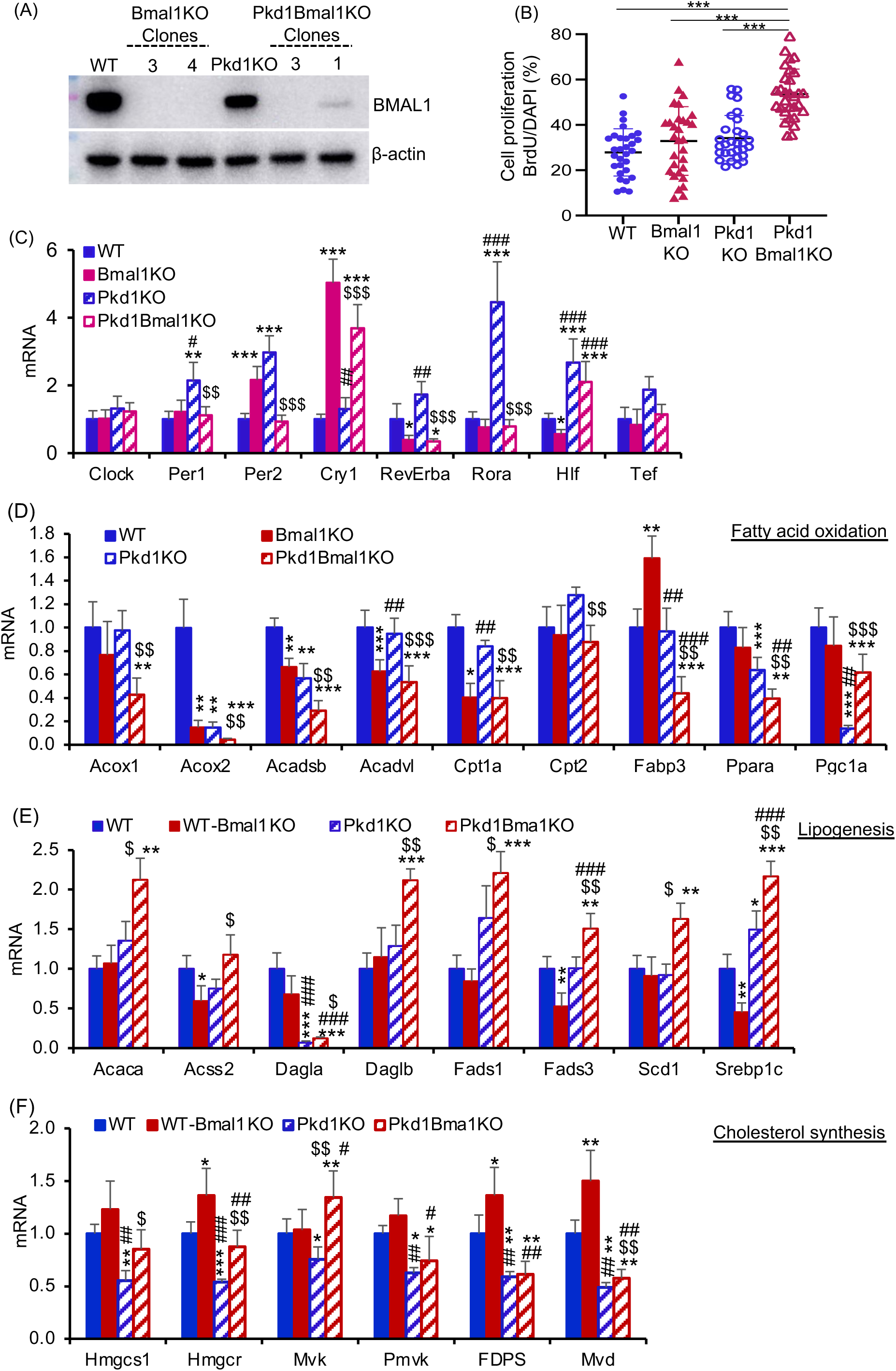
*Bmal1* gene knockout increased cell proliferation and altered the expression of clock genes and lipid metabolism-related genes in *Pkd1*KO mIMCD3 cells. A) Western blot shows WT and *Bmal1*KO clones in WT and *Pkd1* gene knockout mIMCD3 cells. WT-*Bmal1*KO clone #4 and *Pkd1Bmal1*KO clone# 3 were used for further studies. (B) Cell proliferation assessed by BrdU incorporation assay. ***P<0.001 by One-way ANOVA. (C) QRTPCR analysis for mRNA levels of clock genes relative to 18S mRNA in cells. (D) mRNA levels of genes that regulate fatty acid oxidation, (E) lipogenesis, and (F) cholesterol synthesis relative to 18S mRNA. * = *vs* WT, # = *vs Bmal1*KO and $ = *vs Pkd1*KO. *P<0.05, **P<0.01, ***P<0.001 by One-way ANOVA. n=4 in C,D,E and F.

We also examined BMAL1’s prospective gene targets. BMAL1, a transcription factor binds to E-box elements on gene promoters ^31^. The *Pkd1* gene promoter contains an E-box element ^32^, although it is unknown if *Bmal1* is a transcriptional regulator of *Pkd1*. Contrary to our expectation, the *Bmal1*KO cells had significantly increased *Pkd1* mRNA levels when compared to WT cells (Supplemental Fig 4A). We also examined gene expression of *Ets1*, *Sp1*, *Rxra*, *Cebpa*, *Hnf4a*, *Fli1*, *Hnf1a*, *Hnf1b*, *Hnf4a*, *Pax8* and *Tead4,* based on a previous Chip-seq analysis ^33^ that identified these genes as targets of BMAL1 in WT mouse kidneys ^9^. Of the genes detected, *Cebpa, Ets1* and *Hnf1b* were significantly reduced in DKO kidneys and *Pkd1Bmal1*KO cells when compared to RC/RC kidneys and *Pkd1*KO cells respectively (Supplemental Fig 4A,B).

### 5. *Bmal1* gene knockout altered the lipid metabolism-related gene expression in *Pkd1Bmal1*KO mIMCD3 cells and DKO mouse kidneys

Reduced fatty-acid oxidation and increased fatty acid synthesis enhance cyst growth in ADPKD ^34-36^. Since *Bmal1* is known to regulate fatty-acid oxidation, lipogenesis and cholesterol metabolism ^37-41^, we examined genes regulating these processes. In *Pkd1Bmal1*KO cells, mRNA levels of fatty-acid oxidation-related genes, *Acox1*, *Acox2*, *Acadsb*, *Acadvl*, *Cpt1a*, *Cypt2*, *Fabp3*, and *Ppara* showed significant decreases, while *Pgc1a* was increased compared to *Pkd1*KO cells (Fig 5D). *Pkd1*KO cells showed significantly reduced *Acox2*, *Ppara* and *Pgc1a* mRNA levels compared to WT cells (Fig 5D). *Acox2*, *Acadsb*, *Acadvl* and *Cpt1a* are common genes that were significantly decreased in both *Bmal1*KO and *Pkd1Bmal1*KO cells compared to WT and Pkd1KO cells respectively (Fig 5D).

Lipogenesis associated genes, *Acaca*, *Acss2*, *Dagla*, *Daglb*, *Fads1*, *Fads3*, *Scd1*, and Srebp1c were significantly increased in *Pkd1Bmal1*KO cells compared to *Pkd1*KO cells (Fig 5E). *Acss2*, *Fads3* and *Srebp1c* are common genes that were upregulated in *Pkd1Bmal1*KO cells, but downregulated in the *Bmal1*KO cells compared to WT and *Pkd1*KO cells respectively (Fig 5E). Compared to WT cells, *Pkd1*KO cells showed significantly increased *Srebp1c*, and reduced *Dagla* mRNA levels (Fig 5E). Genes related to cholesterol synthesis including *Hmgcs1*, *Hmgcr*, *Mvk* and *Mvd* were significantly increased in *Pkd1Bmal1*KO cells when compared to *Pkd1*KO cells (Fig 5F). When compared to WT cells, the *Pkd1*KO cells showed significantly reduced levels of *Hmgcs1*, *Hmgcsr*, *Mvk*, *Pmvk*, *Fdps* and *Mvd* (Fig 5F). Other fatty acid oxidation and lipogenesis-related genes that remained unchanged are shown in Supplemental Fig 5A,B.

Examination of lipogenesis-related genes in our mouse study showed significantly increased mRNA levels of *Acaca*, *Dagla*, *Fads1*, *Scd1*, *Srebp1c*, *Mag1* and *Lpcat* in DKO mouse kidneys compared to RC/RC kidneys (Fig 6A-G). Furthermore, tissue triglyceride levels showed a 2-fold increase in DKO mouse kidneys when compared to RC/RC mouse kidneys (Figure 6H). To determine if elevated lipogenesis contributes to increased cell proliferation, we inhibited lipogenesis in Pkd1Bmal1KO cells. Firsocostat (GS-0976) is a pharmacological inhibitor of acetyl-CoA carboxylase (ACACA & ACACB), a lipogenesis regulating enzyme ^42^. We found ∼2 fold increase in *Acaca* gene in both DKO mouse kidneys and in Pkd1Bmal1KO cells when compared to RC/RC mouse and Pkd1KO cells respectively (Fig 5E & 6A). Firsocostat (10nM) treatment for 24h significantly reduced cell proliferation in Pkd1Bmal1KO cells, indicated by significant reduction in BrdU incorporation compared to vehicle treated cells, while Pkd1KO cells remained unaffected (Fig 6I). Upon extended Firsocostat treatment for 48h, cell viability indicated by MTT assay was significantly reduced in both Pkd1KO and Pkd1Bmal1KO cells (Figure 6J). The results suggest that the increased cell proliferation in Pkd1Bmal1KO cells can be rescued by inhibition of lipogenesis.

**Fig 6.**
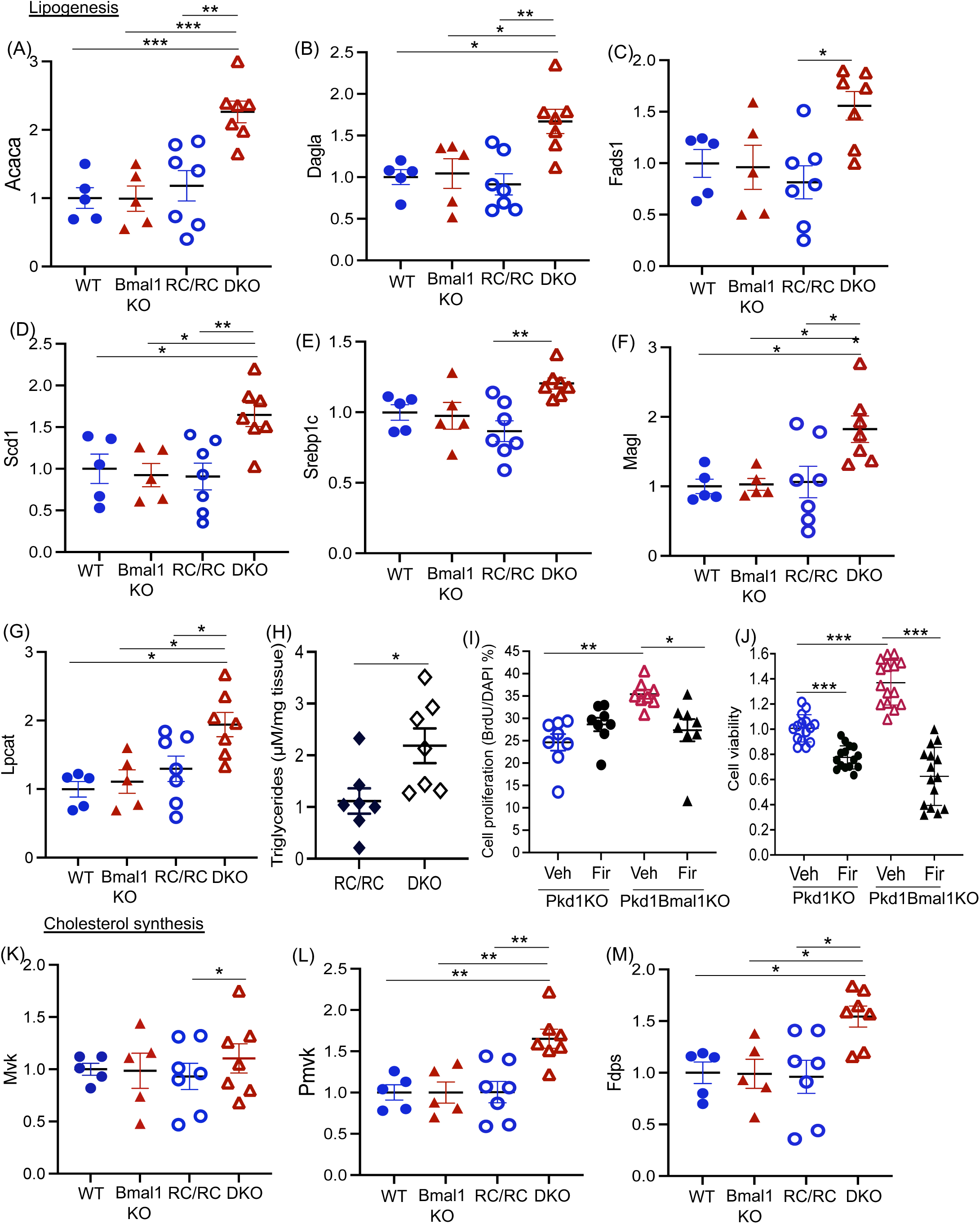
Increased lipogenesis in Bmal1 gene deletion increases lipogenesis in RC/RC mouse kidneys and in Pkd1KO cells. The mRNA levels of genes that regulate lipogenesis, (A) *Acaca*, (B) *Dagla*, (C) *Fads1*, (D) *Scd1*, (E) *Srebp1c*, (F) *Magl*, (G) *Lpcat*, relative to 18S. (H) Mouse kidney tissue triglyceride levels. (I) Pkd1KO and Pkd1Bmal1KO IMCD3 cells were treated with Firsocostat (Fir), (10nM) or vehicle for 24h. Cell proliferation (% BrDU/ DAPI) shown. (J) MTT assay for cell viability of Pkd1KO and Pkd1Bmal1KO IMCD3 cells treated with Firsocostat (10nM) or vehicle for 48h. (K) Cholesterol metabolism related genes *Mvk*, (L) *Pmvk,* and (M) *Fdps*. *P<0.05, **P<0.01, ***P<0.001 by One-way ANOVA.

Cholesterol metabolism-related *Mvk*, *Pmvk* and *Fdps6* genes were also significantly increased in DKO kidneys compared to RC/RC kidneys (Fig 6K,L,M). Fatty-acid oxidation-related genes were not significantly different between DKO and RC/RC kidneys, although RC/RC kidneys showed significantly reduced expression of *Acox1*, *Acadsb*, *Ppara*, *Acat1*, *Acat2* and *Acads* compared to WT kidneys (Supplemental Fig 6 A-N). Unchanged lipogenesis and cholesterol synthesis-related genes in mouse kidneys are shown in Supplemental Fig 7A-J

## Discussion

Our studies show that disruption of the circadian clock stimulates accelerated cyst growth in ADPKD. We found reduced *Bmal*1 gene expression in human nephrectomy ADPKD kidneys, and in *Pkd1* gene knockout mIMCD3 and proximal tubule cells, but not in *Pkd1* gene hypomorphic RC/RC mouse kidneys. Renal collecting duct specific gene deletion of *Bmal1* in RC/RC male mice increased renal cell proliferation, apoptosis, fibrosis, and cyst growth. *Bmal1* gene deletion in RC/RC mouse kidneys significantly altered the expression of other clock genes, reduced the expression of BMAL1 target genes and increased lipogenesis. *In vitro,* proliferation of *Pkd1Bmal1*KO mIMCD3 cells was higher than *Pkd1*KO mIMCD3 cells, which could be rescued by inhibition of lipogenesis.

Our finding that *Bmal1* gene knockout in the renal collecting ducts increased cyst growth in ADPKD mice is significant because it shows for the first time that chronodisruption accelerates ADPKD progression. Chronodisruption is disease inducing, and known to directly drive metabolic dysfunction, cause type-2 diabetes, obesity, various cancers and neurodegenerative conditions, and adversely affect fetal development ^43-47^. Since chronodisruption by alterations in behavior or the environment are systemic effects, we used renal collecting duct specific *Bmal1* gene deletion as a model for genetic disruption of the molecular clock so as to focus on the kidney. In our study, *Bmal1* gene knockout altered the expression of clock genes including *Cry1*, *Per1*, *Per2*, and *RevErba* that are in the negative arm of the clock, and are known to be regulated by *Bmal1*.

We found that collecting duct-specific gene deletion of *Bmal1* dramatically increased cyst growth, cell proliferation and fibrosis in the male RC/RC mouse model of ADPKD. In prior studies, *Bmal1* gene knockout reduced locomotor activity, and induced tendon calcification, sarcopenia, hypersensitivity to insulin-shock, fragmented sleep, and rapid ageing in mice ^48, 49^. In the kidneys, nephron-specific *Bmal1* gene deletion caused Na^+^ retention in response to a K^+^- restricted diet ^50^, lowered blood pressure ^50-52^ and altered the urinary metabolome in mice ^48^. However, the effect of altering *Bmal1* gene expression on kidney disease progression is context dependent, as shown in experimental mouse models ^53^. In a mouse model of AKI, shRNA mediated *Bmal1* gene knockout alleviated AKI, while adenoviral overexpression of *Bmal1* accelerated renal injury ^54^. Similarly in unilateral ureteral obstruction model of CKD, global gene knockout of *Bmal1* reduced fibrosis ^55^ or had no effect on fibrosis or disease progression ^56^. In contrast to the above mentioned studies, and supporting our results, renal tubule specific *Bmal1* gene knockout worsened hyperglycemia, and accelerated renal hypertrophy in mice with streptozotocin-induced type-1 diabetes, compared to similarly treated WT mice ^57^. Moreover, global or proximal tubule specific gene knockout of *Bmal1* worsened disease progression in the adenine induced model of CKD in mice ^58,59^. We found that in the presence of *Pkd1* gene mutation, *Bmal1* gene knockout in renal collecting ducts increased cell proliferation indicated by increase in Ki-67 immunostaining, and pERK/ERK1/2, pCREB/CREB, pS6/S6 and CyclinD1 levels, all factors known to increase cell proliferation in ADPKD kidneys ^25^. Our data is consistent with previous studies where *Bmal1* gene deletion or inactivation increases the proliferation rate of cancer cells ^60, 61^.

BMAL1 binds to E-box elements on promoters of rhythmically and non-rhythmically expressed genes ^31^. Over 7 million E-box elements exist in the mouse genome, ∼ 0.7% of which are bound by CLOCK-BMAL1 or NPAS2-BMAL1 dimers in mouse peripheral tissues ^33, 62^. Based on this published data, we examined 12 prospective gene targets of BMAL1 and identified *Ets1*, *Hnf1b* and *Cebpa* to be significantly reduced in *Bmal1* deficient *Pkd1*KO cells and RC/RC mouse kidneys. C/EBPα plays an essential role in embryonic development and supresses cell proliferation by stabilizing p21 and inhibiting *cdk2*, *cdk4* and *E2F* ^63^. Inhibiting C/EBPα contributes to tumor progression in liver, pancreatic, breast and lung cancers ^63^. Similarly, ETS1 and HNF1 transcription factors have oncogenic or tumor supressor roles in a context-dependent manner in human cancers ^64, 65^. HNF1β is known to regulate *Pkd2* and *Pkhd1* gene expression, while and ETS1 regulates *Pkd1* gene expression ^32, 66, 67^.

Fatty-acid metabolism is impaired in ADPKD and is thought to enhance renal cyst growth ^34-36^. A previous study showed reduced fatty-acid oxidation-related gene expression in ADPKD mouse kidneys, and treatment with fenofibrate, a PPARα agonist inhibited cyst growth in these mice ^34^. Similarly, pups born from female mice fed with a high fat diet had higher cystic index compared to those fed with low fat diet ^35^. *In vitro* studies have also shown that *Pkd1* gene mutation leads to defects in fatty-acid oxidation ^35, 36^. Consistent with these studies, we found fatty-acid oxidation-related genes to be significantly reduced in *Pkd1*KO cells compared to WT cells, and in RC/RC mouse kidneys compared to WT mouse kidneys. BMAL1 is known to regulate fatty-acid oxidation ^37-41^. Bignon *et al* ^41^ showed that renal tubule specific *Bmal1* gene deletion in WT mice significantly reduced mRNA and protein expression of ACADS, ACADM, ACADL, and ACADVL which catalyze the rate-limiting step of β-oxidation, suggesting disrupted fatty-acid oxidation. However in our study, renal collecting duct specific *Bmal1* gene deletion in the *Bmal1*KO mice or DKO mice did not alter fatty-acid oxidation-related genes when compared to WT or RC/RC mouse kidneys. The reason for this disparity is presently unclear. Our results from the mIMCD3 cells were different, with 4 of the 14 fatty acid oxidation-related genes showing significant decreases in the *Bmal1*KO and *Pkd1Bmal1*KO cells compared to WT and *Pkd1*KO cells respectively.

Unlike fatty-acid oxidation, our *in vitro* and *in vivo* results consistently showed increased lipogenesis and cholesterol metabolism-related genes in *Bmal1* gene deleted cells and mouse kidneys. The mRNA levels of 7 out of the 14 lipogenesis-related genes tested in *Pkd1Bmal1*KO cells and in DKO kidneys showed significant increase compared to WT cells and RC/RC kidneys. Of the lipogenesis-related genes, *Acaca*, *Fads1*, *Scd1* and *Srebp1c* were upregulated in both DKO mouse kidneys and in *Pkd1Bmal1*KO cells, suggesting regulation by *Bmal1.* Tissue triglyceride levels were also higher in DKO mouse kidneys compared to RC/RC mouse kidneys, supporting increased lipogenesis. Importantly, pharmachological inhibition of lipogenesis rescued the increased cell proliferation in *Pkd1Bmal1*KO cells. The *Pkd1Bmal1*KO cells and DKO kidneys showed significant increases in several cholesterol synthesis-related genes. Mvk was a frequently upregulated genes in the cells and kidneys. Renal collecting duct specific *Bmal1* gene deletion in WT mice (*Bmal1*KO mice) did not alter any lipogenesis, cholesterol synthesis or fatty-acid oxidation-related genes when compared to WT mouse kidneys, suggesting that *Bmal1* gene knockout in the ADPKD kidneys is important for these changes.

Although *Bmal1* gene deletion altered the expression of multiple clock genes, BMAL1 regulated genes and lipid metabolism-related genes in the mIMCD3 cells and mouse kidneys, their changes were not always similar in the cells and kidneys. This could be because *Bmal1* is deleted only in the collecting duct cells of the mouse kidneys, while the IMCD3 cells had complete gene deletion of *Bmal1*. Moreover, the mice were all sacrificed around 12 PM, while the mIMCD3 cells were collected 24h after synchronization *in vitro*.

While patients with ADPKD often develop non-dipping hypertension, polyuria, polydipsia, and nocturnal polyuria ^68-70^, it is not known if chronodisruption is a modifier of ADPKD progression. Our results show that *Bmal1* gene knockout in the renal collecting ducts disrupt the circadian clock and expression of key elements of cell proliferation, fibrosis and lipid metabolism, thereby accelerating cyst growth in ADPKD. It is possible, therefore, that chronodisruption in individuals with ADPKD might be a cause of, or contribute to more rapid disease progression. Environmental or behavioral factors such as mistimed feeding, shift work or jet lag are known to induce chronodisruption ^71^. Shift work for instance, reduces the estimated GFR in humans and is a risk factor for CKD ^72-74^, cardiovascular disease ^75^, multiple cancers ^76^ and metabolic disorders ^77^. It is unclear if environmental or behavioral factors have any effect on ADPKD progression. Future studies could examine if poor quality or duration of sleep or mistimed eating habits act as chronodisruptors and accelerate ADPKD progression, and if methods to re-set the clock using pharmacological approaches, or behavioral modifications can slow progression of ADPKD.

## Acknowledgments

We thank Dr. Peter Harris and Dr. Katharina Hopp for providing *Pkd1*^RC/RC^ mice; Dr. Stefan Somlo and Dr. Yiqiang Cai (George M. O’Brien Kidney Center, Yale University, New Haven, CT) for *Pkd1*KO and control proximal tubular and inner medullary collecting ducts cells; Dr. Peter Igarashi for the Pkhd1^cre^ mice and Dr. Darren Wallace for human kidney tissues (NIDDK PKD-Research Resource Consortium and the Kansas PKD Research and Translational Core Center-U54 DK126126).

## Author contributions

RR conceptualized and designed studies, analyzed the results, and wrote the paper. AJ performed experiments, analyzed results and wrote parts of the paper. CJW, VR, MMS and NSP performed some studies, analyzed results and edited the paper. MLG conceptualized some studies and reviewed the paper. All authors read, edited and approved the paper.

## Data sharing statements

No proteomics, transcriptomics or GWAS data was generated in this paper. All in vivo studies using mouse models were done, and tissues analyzed at the University of Kansas Medical Center. All data that was generated are provided in the figures in the main text or in Supplementary Material. The datasets used to examine Bmal1 expression levels in human samples are openly available on the kidney interactive transcriptomics database (https://humphreyslab.com/SingleCell/) and permission was obtained to use it. If any added information is required to re-examine or analyze the data shown in this paper, they will be readily available from the lead author. All requests for data will be processed based on institutional policies for noncommercial research purposes. Data sharing could require a data transfer agreement as determined by the University of Kansas Medical Center’s legal department.

