## Supplemental materials for "Circadian clock disruption and growth of kidney cysts in autosomal dominant polycystic kidney disease"

**Supplemental Figure 1. *Bmal1* expression in WT and RC/RC mouse kidneys**

**Supplemental Figure 2. Cell proliferation in mouse kidneys**

**Supplement Figure 3. *Bmal1* gene knockout clones of WT and *Pkd1*KO mIMCD3 cells**

**Supplemental Figure 4. BMAL1 regulated genes**

**Supplemental Figure 5. Fatty acid oxidation and lipogenesis-related gene expression in *Pkd1*KO mIMCD3 cells.**

**Supplemental Figure 6. Fatty-acid oxidation-related gene expression in mouse kidneys.**

**Supplemental Figure 7. Lipogenesis and cholesterol metabolism-related gene expression in mouse kidneys**

**Supplemental Table 1: Primer list**

**Supplemental methods**

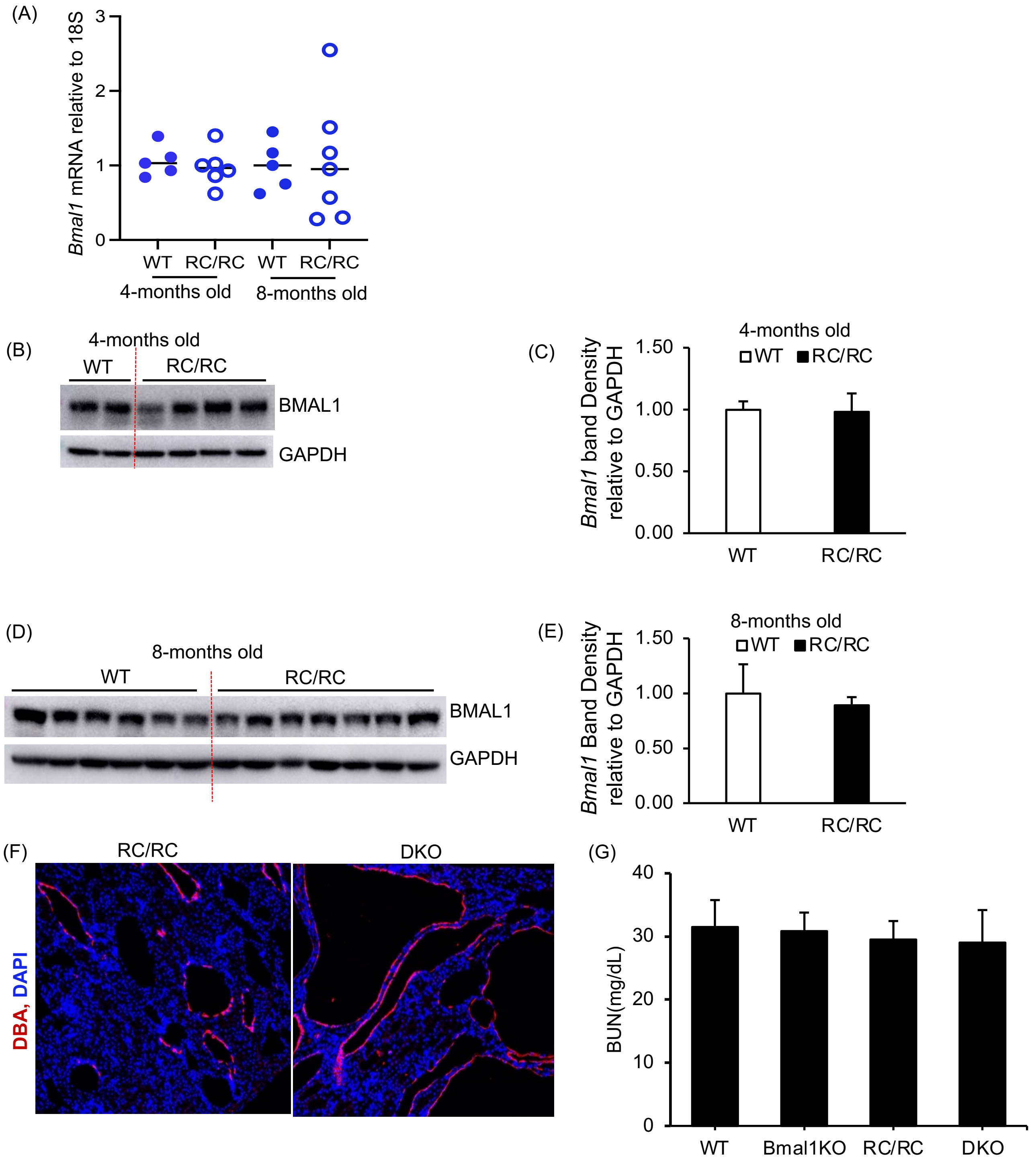
(A)
Bmal1 mRNA relative to 18S
0
1
2
3
WT
RC/RC
WT
RC/RC
4-months old
8-months old
(B)
4-months old
WT
RC/RC
BMAL1
GAPDH
(C)
4-months old
WT
RC/RC
Bmal1 band Density relative to GAPDH
0.00
0.50
1.00
1.50
WT
RC/RC
(D)
8-months old
WT
RC/RC
BMAL1
GAPDH
(E)
8-months old
WT
RC/RC
Bmal1 Band Density relative to GAPDH
0.00
0.50
1.00
1.50
WT
RC/RC
(F)
RC/RC
DKO
DBA, DAPI
(G)
BUN(mg/dL)
0
10
20
30
40
WT
Bmal1KO
RC/RC
DKO

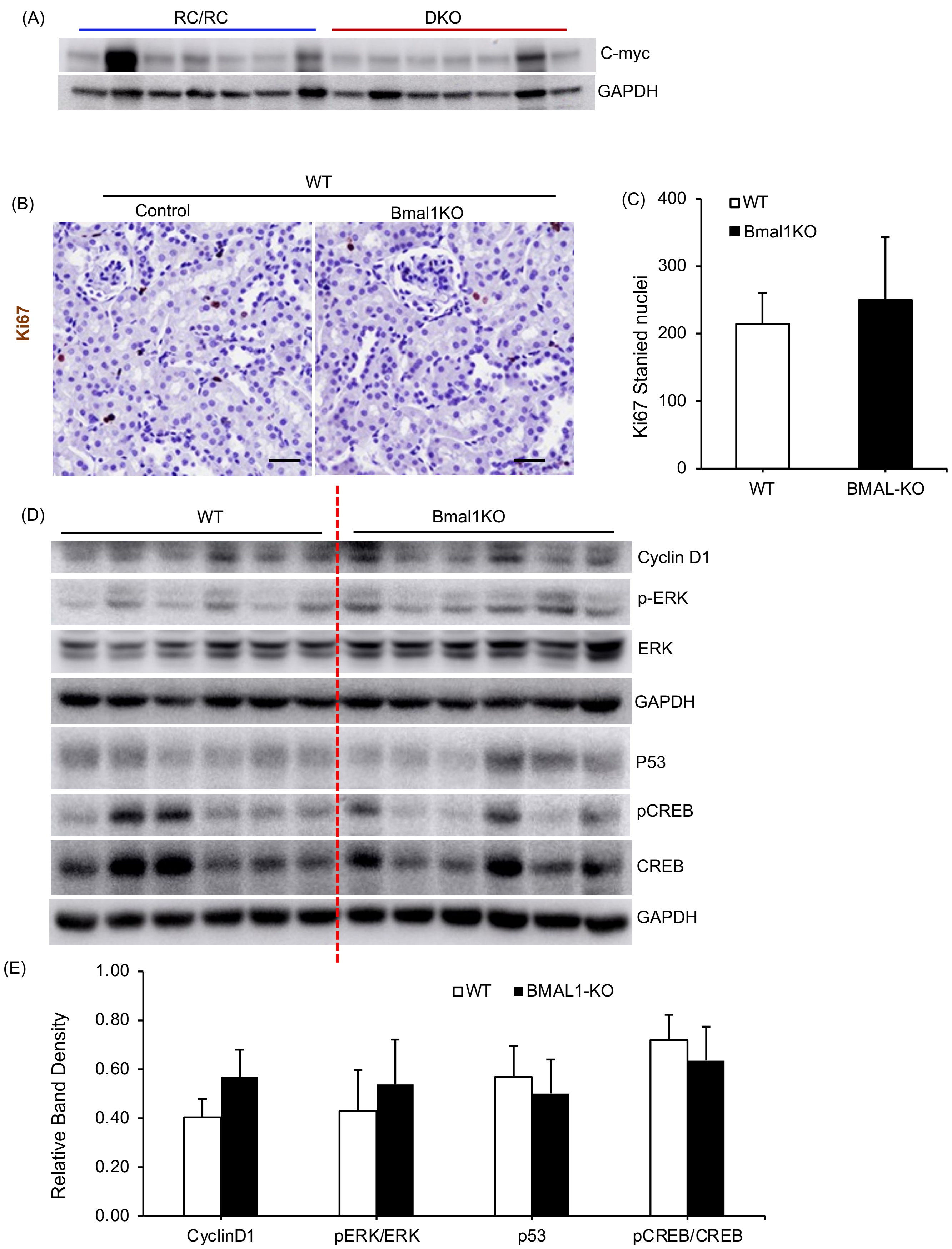
(A)
RC/RC
DKO
C-myc
GAPDH
(B)
WT
Control
Bmal1KO
Ki67
(C)
400
300
200
100
0
WT
Bmal1KO
Ki67 Stanied nuclei
WT
BMAL-KO
(D)
WT
Bmal1KO
Cyclin D1
p-ERK
ERK
GAPDH
P53
pCREB
CREB
GAPDH
(E)
1.00
0.80
0.60
0.40
0.20
0.00
WT
BMAL1-KO
Relative Band Density
CyclinD1
pERK/ERK
p53
pCREB/CREB

(A) **WT_BMAL1_4**

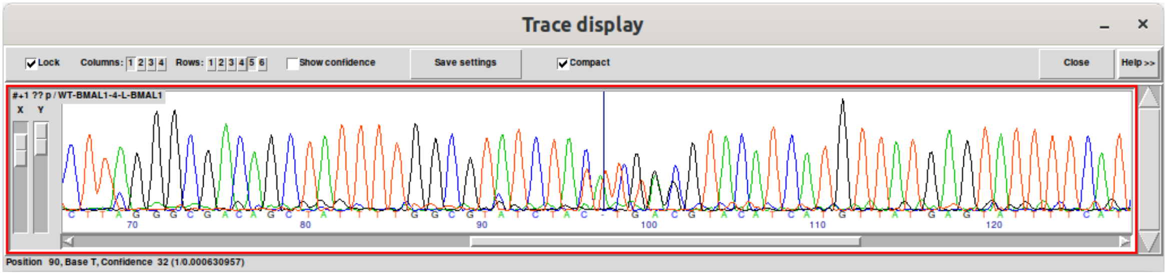


```
GGCGACAGCTATTTTGGCGTATCTAC---TTGAC-------GTACATCATGTTATGAGTATTTTCATCA
||||||||||||||||||||||||||               ||||||||||||||||||||||||||||
GGCGACAGCTATTTTGGCGTATCTACCACAGGAACTTCTAGGTACATCATGTTATGAGTATTTTCATCA
  A  T  A  I  L  A  Y  L  P  Q  E  L  L  G  T  S  C  Y  E  Y  F  H
  A  T  A  I  L  A  Y  L     L  D         V  H  H  V  M  S  I  F  IKTT+
```

(B) **PKD1_KO_BMAL1_3**

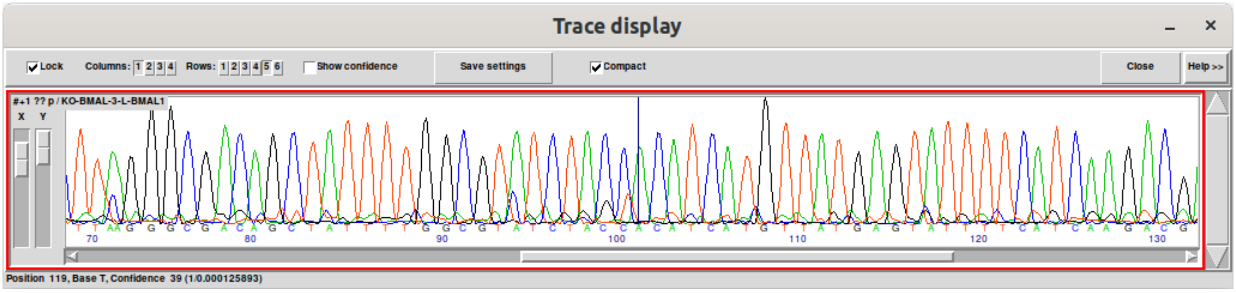


```
GGCGACAGCTATTTTGGCGTATCTACC----------------ACATCATGTTATGAGTATTTTCATCA
|||||||||||||||||||||||||||||||||||||||||||||||||||||||||||||||||||||
GGCGACAGCTATTTTGGCGTATCTACCACAGGAACTTCTAGGTACATCATGTTATGAGTATTTTCATCA
  A  T  A  I  L  A  Y  L  P  Q  E  L  L  G  T  S  C  Y  E  Y  F  H
  A  T  A  I  L  A  Y  L  P                  H  H  V  M  S  I  F  IKTT+
```

(A)

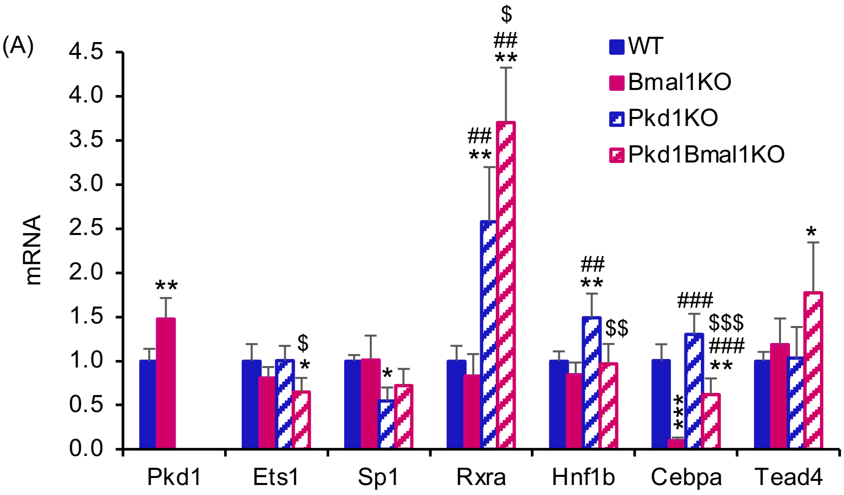
WT
Bmal1KO
Pkd1KO
Pkd1Bmal1KO
mRNA
0.0
0.5
1.0
1.5
2.0
2.5
3.0
3.5
4.0
4.5
Pkd1
Ets1
Sp1
Rxra
Hnf1b
Cebpa
Tead4

(B)

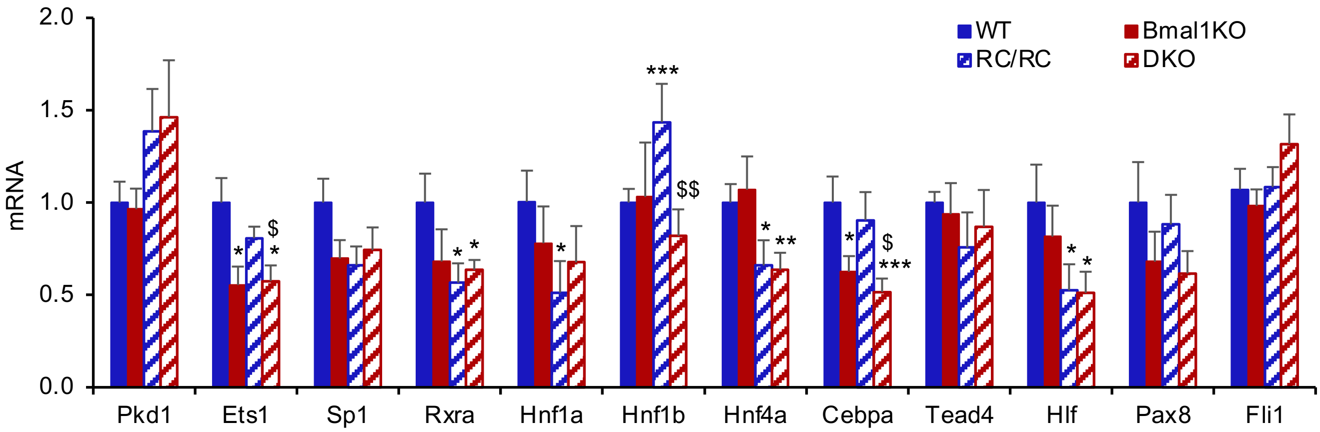
WT
Bmal1KO
RC/RC
DKO
mRNA
0.0
0.5
1.0
1.5
2.0
Pkd1
Ets1
Sp1
Rxra
Hnf1a
Hnf1b
Hnf4a
Cebpa
Tead4
Hlf
Pax8
Fli1

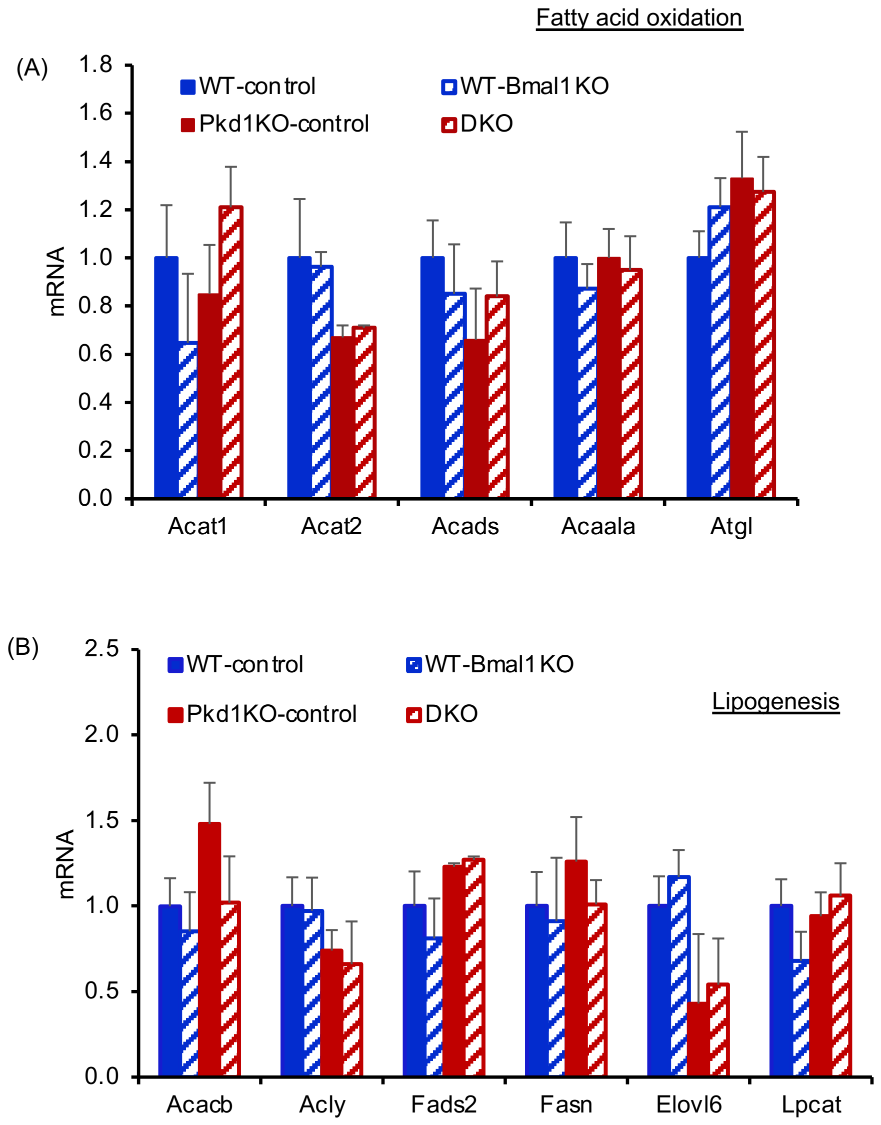
Fatty acid oxidation
(A)
1.8
1.6
1.4
1.2
1.0
0.8
0.6
0.4
0.2
0.0
mRNA
WT-control
WT-Bmal1 KO
Pkd1KO-control
DKO
Acat1
Acat2
Acads
Acaala
Atgl
(B)
2.5
2.0
1.5
1.0
0.5
0.0
mRNA
WT-control
WT-Bmal1 KO
Pkd1KO-control
DKO
Lipogenesis
Acacb
Acly
Fads2
Fasn
Elovl6
Lpcat

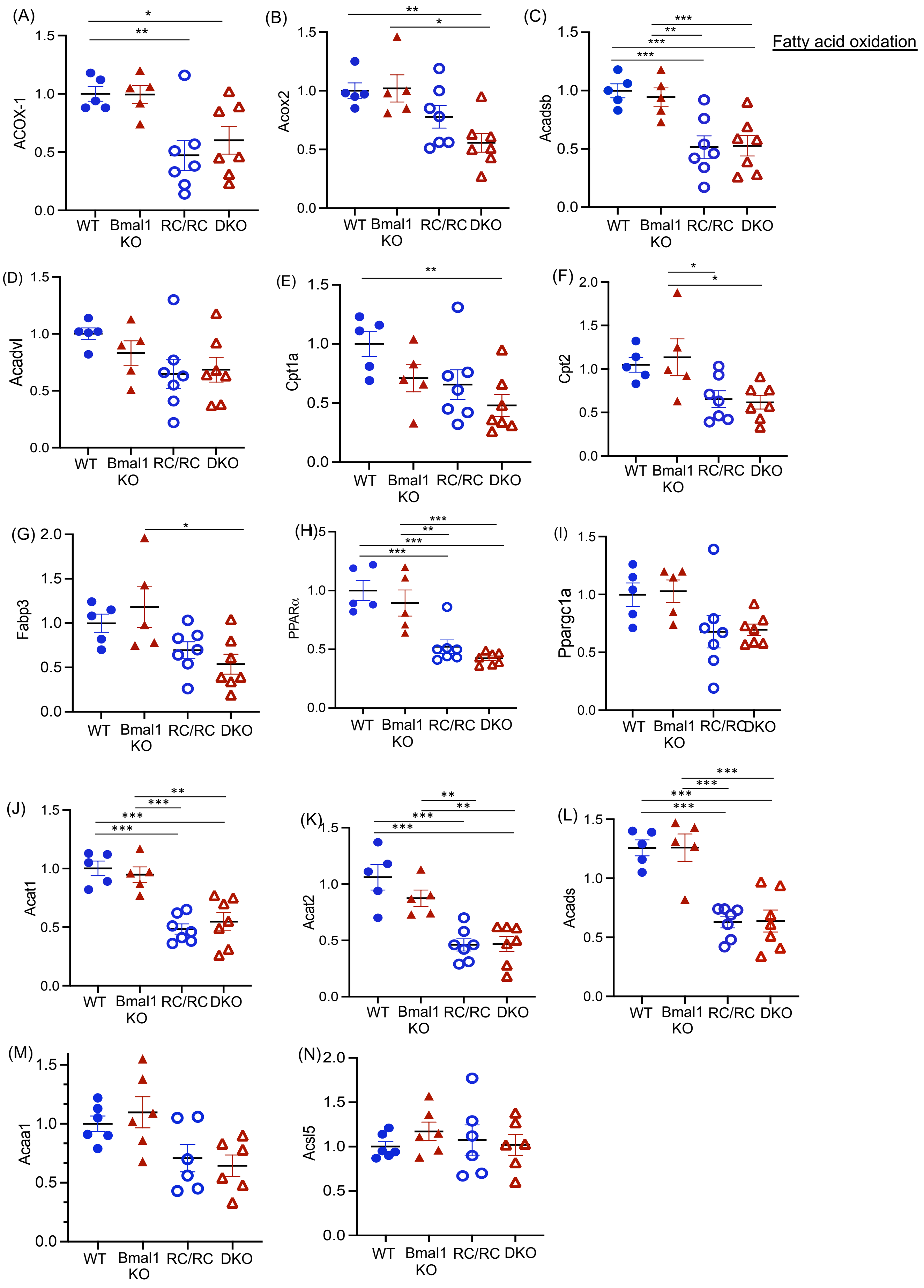
Fatty acid oxidation
(A)
ACOX-1
(B)
Acox2
(C)
Acadsb
(D)
Acadvl
(E)
Cpt1a
(F)
Cpt2
(G)
Fabp3
(H)
PPARα
(I)
Ppargc1a
(J)
Acat1
(K)
Acat2
(L)
Acads
(M)
Acaa1
(N)
Acsl5
WT
Bmal1 KO
RC/RC
DKO

Lipogenesis

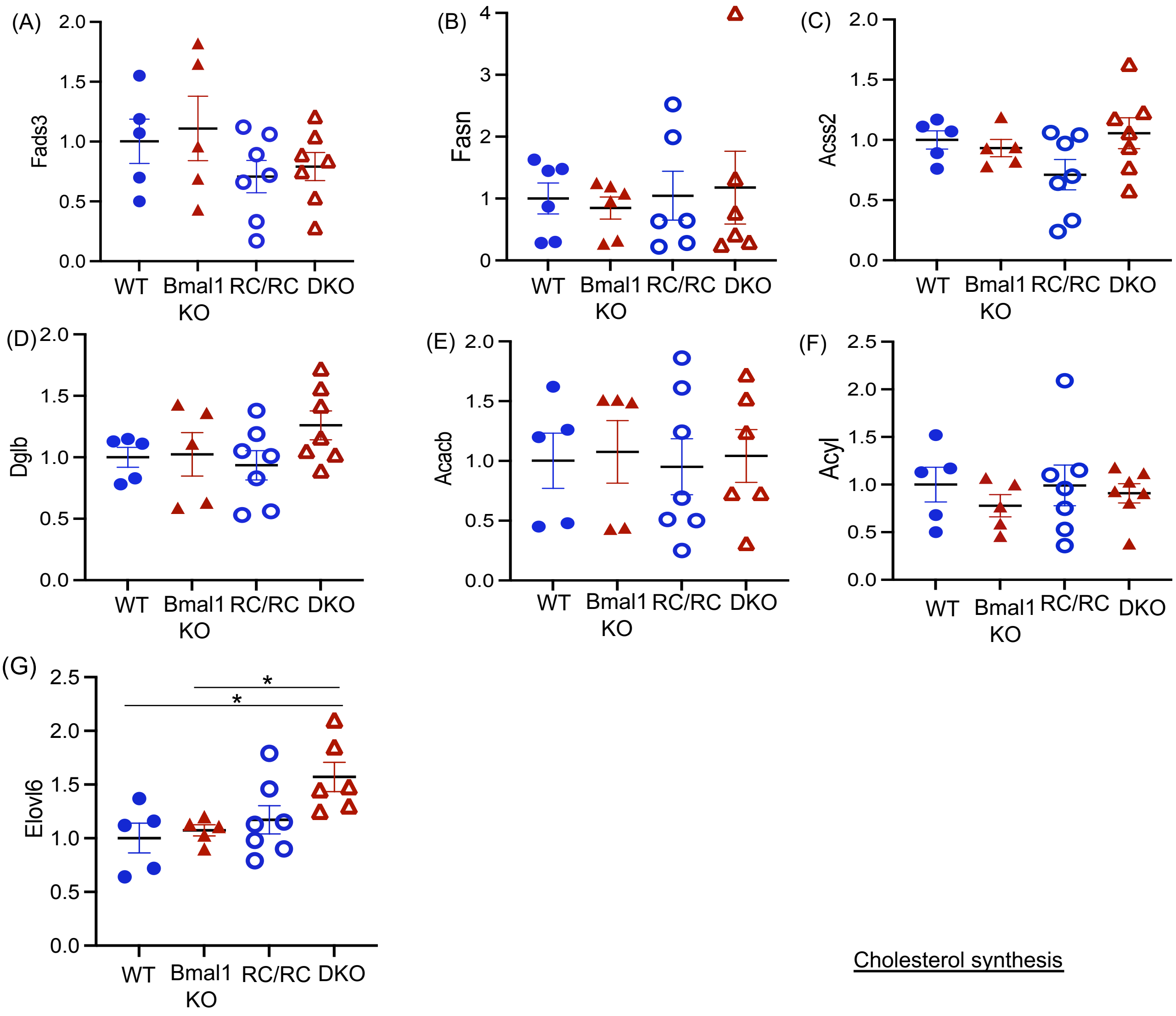


Cholesterol synthesis

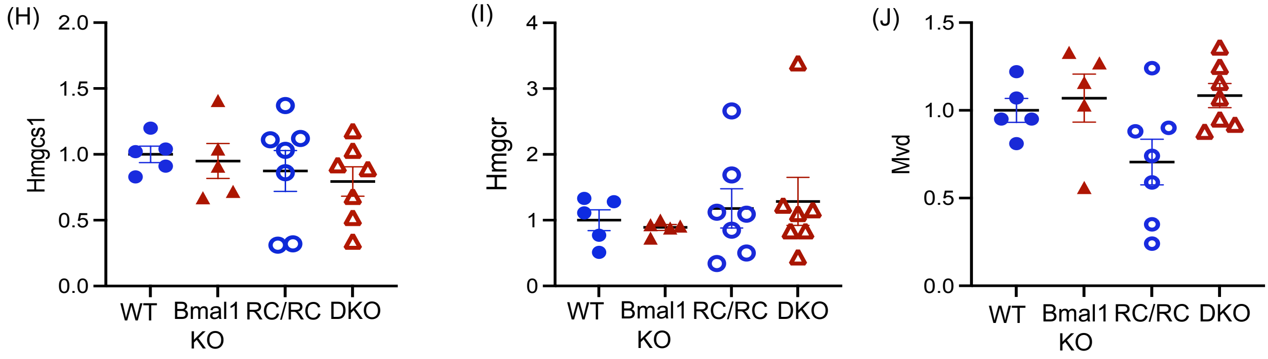

**Supplemental Figure legends:**

**Supplemental 1. *Bmal1* expression in WT and RC/RC mouse kidneys:** (A) *Bmal1* mRNA expression relative to 18S mRNA measured by QRTPCR in WT and RC/RC kidneys at 4 and 8 months of age. (B) Western blot for BMAL1 and (C) densitometry for 4 month old mouse kidneys (WT, n=2, RC/RC, n=4), and (D) Western blot and (E) densitometry for 8 month old mouse kidneys (WT, n=6, RC/RC, n=7). (F) Immunostaining for DBA (red) and DAPI (blue) (G) BUN assay.

**Supplemental 2. Cell proliferation in mouse kidneys:** (A) Western blot for c-Myc. (B) Ki67 stained kidney section (Scale bar 100μm) and (C) Ki-67 quantitation (n=6) (D) Western blot and (E) densitometry for pro-proliferative proteins in WT and *Bmal1*KO mouse kidneys (n=6)..

**Supplement 3. *Bmal1* gene knockout clones of WT and *Pkd1*KO mIMCD3 cells:** (A) Gene knockout of *Bmal1* by CRISPR/Cas12a in WT and (B) *Pkd1* gene knockout mIMCD3 cells. Sequences of wild-type mIMCD3 (WT), and the selected clones #4 of WT and clone #3 of *Pkd1*KO mIMCD3 cells, validating the deletion of base pairs in exon 2.

**Supplemental 4. BMAL1 regulated genes:** (A) QRTPCR for mRNA levels of BMAL1 target genes relative to 18S mRNA in mIMCD3 cell lysates and (B) in mouse kidney tissue lysates. * = *vs* WT, # = *vs* *Bmal1*KO and $ = *vs* *Pkd1*KO. *P<0.05, **P<0.01, ***P<0.001 by by One-way ANOVA.

**Supplemental 5. Fatty acid oxidation and lipogenesis-related gene expression in *Pkd1*KO mIMCD3 cells.** (A) QRTPCR analysis for mRNA levels of genes related to fatty acid oxidation (n=4) and (B) lipogenesis (n=4) relative to 18S mRNA.

**Supplemental 6. Fatty-acid oxidation-related gene expression in mouse kidneys.** (A-N) The mRNA levels of genes that regulate fatty-acid oxidation relative to 18S mRNA. *P<0.05, **P<0.01, ***P<0.001 by One-way ANOVA.

**Supplemental 7. Lipogenesis and cholesterol metabolism-related gene expression in mouse kidneys.** (A-G) QRTPCR analysis for mRNA levels of genes that regulate lipogenesis and (H-J) cholesterol metabolism relative to 18S mRNA in whole kidney lysates. . *P<0.05, by One-way ANOVA.

**Supplemental Table-1**
**Mouse primers sequence used or QRTPCR**

| | Gene Name | Primer Forward (F) and Reverse (R) |
|---|---|---|
| 1 | αSMA | F - TCAGGGAGTAATGGTTGGAATG<br>R - GGTGATGATGCCGTGTTCTA |
| 2 | Collagen-1a | F - AGACATGTTCAGCTTTGTGGAC<br>R - GCAGCTGACTTCAGGGATG |
| 3 | Collagen-3a | F - TCCCCTGGAATCTGTGAATC<br>R - TGAGTCGAATTGGGGAGAAT |
| 4 | Fibronectin | F - ATGTGGACCCCTCCTGATAGT<br>R - GCCCAGTGATTTCAGCAAAGG |
| 5 | Kim1 | F - AAACCAGAGATTCCCACACG<br>R - GTCGTGGGTCTTCCTGTAGC |
| 6 | Ngal | F - ATGGCCCTGAGTGTCATGTGT<br>R- GCTCCAGATGCTCCTTGGTATG |
| 7 | Acads | F - CGTGACTTTGCCGAGAAGGAGT<br>R - TCAGCTCCTCTGGCACATCCAT |
| 8 | Acadsb | F - TGGAAGCCACACGGTTGCTAAC<br>R - CATCCACTCGATGCACTTGCTTG |
| 9 | Acadvl | F - GAGAAGGTGGAGGACGACACTT<br>R - CACAATCTCTGCCAAGCGAGCA |
| 10 | Acss2 | F - AGGTGACCAAGTTCTACACGGC<br>R - GTTGATGGGTTCACCTACTGTGC |
| 11 | Acox1 | F - CTTGGATGGTAGTCCGGAGA<br>R - TGGCTTCGAGTGAGGAAGTT |
| 12 | Acox2 | F - TACCAACGCCTGTTTGAGTG<br>R - TTTCCAGCTTTGCATCAGTG |
| 14 | Cpt2 | F - CAATGAGGAAACCCTGAGGA<br>R - GATCCTTCATCGGGAAGTCA |
| 15 | Aacat2 | F - GAGATTGTGCCAGTGCTGGTGT<br>R - GTGACAGTTCCTGTCCCATCAG |
| 16 | Acat1 | F - GCAGGGAAGTTTGCCAGTGAGA<br>R - GAACACGGTCTTGAGCTTTGGC |
| 17 | Ppara | F - ACCACTACGGAGTTCACGCATG<br>R - GAATCTTGCAGCTCCGATCACAC |
| 18 | Pgc1a | F - GAATCAAGCCACTACAGACACCG<br>R - CATCCCTCTTGAGCCTTTCGTG |
| 19 | Cpt1a | F - GGCATAAACGCAGAGCATTCCTG<br>R - CAGTGTCCATCCTCTGAGTAGC |
| 20 | Acaa1a | F - ATGACCTCGGAGAATGTGGCTG<br>R - AGGACAGTGGTTGTCACAGGCA |
| 21 | Acly | F - AGGAAGTGCCACCTCCAACAGT<br>R - CGCTCATCACAGATGCTGGTCA |
| 22 | Acaca | F - GTTCTGTTGGACAACGCCTTCAC<br>R - GGAGTCACAGAAGCAGCCCATT |
| 23 | Acacb | F - AGAAGCGAGCACTGCAAGGTTG<br>R - GGAAGATGGACTCCACCTGGTT |
| 24 | Fasn | F - CACAGTGCTCAAAGGACATGCC |

| | | |
|---|---|---|
| | | R - CACCAGGTGTAGTGCCTTCCTC |
| 25 | Scd1 | F - GCAAGCTCTACACCTGCCTCTT<br>R - CGTGCCTTGTAAGTTCTGTGGC |
| 26 | Mvk | F - ATGCTTCAGCGACTGGACACGA<br>R - AGCAGAGCCATGCCTTCATTGC |
| 27 | Magl | F - GACACCATCCAGAAGGACTACC<br>R - GATTGGCAAGGACCAGAGGTGA |
| 28 | Fads1 | F - ACCTGTCAGTCTTTGGCACCTC<br>R - TCCTTGCGGAAGCAGTTAGGCT |
| 29 | Fads2 | F - TTCCTGGAGAGCCACTGGTTTG<br>R - GAAGAAGGACTGCTCCACATTGC |
| 30 | Fads3 | F - TGGCGTACATGCTGGTGTGCAT<br>R - CCTGACAGCAACAAAGAGGAGC |
| 31 | Lpcat | F - CCTGTATGCCAGCAATGTGAGG<br>R - TCAGCAGGCAAGCGAAGTTGTC |
| 32 | Atgl | F - GGAACCAAAGGACCTGATGACC<br>R - ACATCAGGCAGCCACTCCAACA |
| 33 | Dagla | F - CATCTCACCAGCCATGCTGCAT<br>R - GCTCACCATGACCTTTGCAGCA |
| 34 | Daglb | F - CAGTGGAAGGTGACCAGCACAA<br>R - CGACCACAACAGACTCCTTCCT |
| 36 | Srebp1c | F - GGAGCCATGGATTGCACATT<br>R - GGCCCGGGAAGTCACTGT |
| 37 | Elovl6 | F - CGGCATCTGATGAACAAGCGAG<br>R - GTACAGCATGTAAGCACCAGTTC |
| 38 | Hmgcs1 | F - GGAAATGCCAGACCTACAGGTG<br>R - TACTCGGAGAGCATGTCAGGCT |
| 39 | Hmgcr | F - GCTCGTCTACAGAAACTCCACG<br>R - GCTTCAGCAGTGCTTTCTCCGT |
| 40 | Pmvk | F - ACTTCGTGACCGAGAGGCTGAA<br>R - CTTGTAGGTGCTCGCATCCAGA |
| 41 | Mvd | F - CAGCTAGTCCACCGCTTCAACA<br>R - CAAACTCAGCCACAGTGTCCTC |
| 42 | Fdps | F - GGTGGTTCAGTGTCTGCTACGA<br>R - CGCCTCATACAGTGCTTTCACC |
| 43 | Fabp3 | F - AGAGTTCGACGAGGTGACAGCA<br>R - TTGTCTCCTGCCCGTTCCACTT |
| 44 | Clock | F- AGAACTTGGCATTGAAGAGTCTC<br>R- GTCAGACCCAGAATCTTGGCT |
| 45 | Bmal1 | F- GCCCCACCGACCTACTCT<br>R-TGTCTGTGTCCATACTTTCTTGG |
| 46 | Per1 | F- GCTTCGTGGACTTGACACCT<br>R- TGCTTTAGATCGGCAGTGGT |
| 47 | Per2 | F- TCCGAGTATATCGTGAAGAACG<br>R- CAGGATCTTCCCAGAAACCA |
| 48 | Cry-1 | F- ATCGTGCGCATTTCACATAC<br>R- TCCGCCATTGAGTTCTATGAT |
| 49 | Reverb-a | F- CTTCCGTGACCTTTCTCAGC<br>R- CAGCTCCTCCTCGGTAAGTG |
| 50 | Ror-a | F- GTGGAGACAAATCGTCAGGAAT |

| | | |
|---|---|---|
| | | R- TGGTCCGATCAATCAAACAGTTC |
| 51 | Ror-b | F- ACAGGAACCGTTGCCAACACTG<br>R- CTTCTGCACCTCAGCATACAGG |
| 52 | Dbp | F- GAGCCTTCTGCAGGGAAACA<br>R- GCCTTGCGCTCCTTTTCC |
| 53 | Hlf | F- CTCTGAGGAAGAACTGAAGCCAC<br>R- TGCGATCTGGTTCTCCTTCAGC |
| 54 | Tef | F- TCGGAAGCACAGGTTTGCAGAG<br>R- GGAGCGTTTAGCTGCCACATTG |
| 55 | Pkd1 | F- TCAATTGCTCCGGCCGCTG<br>R-CCAGCGTCTGAAGTAGGTTGTGGG |
| 56 | Ets1 | F-CCAGAATCCTGTTACACCTCGG<br>R-CAGCGTCTGATAGGACTCTGTG |
| 57 | Sp1 | F-CTCCAGACCATTAACCTCAGTGC<br>R-CACCACCAGATCCATGAAGACC |
| 58 | RxRa | F-GTGAAAGATGGGATTCTCCTGGC<br>R- GTCACGCATCTTAGACACCAGC |
| 59 | Hnf1a | F- AGAGACCTTGGTGGAGGAGTGT<br>R- GGCAAACCAGTTGTAGACACGC |
| 60 | Hnf1b | F-GCCTTAGTGGAGGAGTGTAACAG<br>R-TCTGCCTGAACGCCTCTTCCTT |
| 61 | Cebpa | F-GCAAAGCCAAGAAGTCGGTGGA<br>R-CCTTCTGTTGCGTCTCCACGTT |
| 62 | Tead4 | F-GCTCTGGATGTTGGAGTTCTCG<br>R-TTGGGCTTGACTGGCTGATGTG |
| 63 | Pax8 | F-TGCTCAGCCTGGCAATGACAAC<br>R-ACGAAGGTGCTTTCGAGGACCA |
| 64 | Fli1 | F- CCATACAGACCAGTCCTCACGA<br>R-CATGGTCTGTGATCCTCCAAGG |
| 65 | 18s | F - GTAACCCGTTGAACCCGA<br>R – CCATCCAATCGGTAGTAGCG |

### Recombination Specificity in *Bmal1*KO and DKO mice

| | Gene Name | Primer Forward (F) and Reverse (R) |
|---|---|---|
| 1 | Mo-Bmal1 | F - ACTGGAAGTAACTTTATCAAACTG<br>R – CTGACCAACTTGCTAACAATTA<br>F- CTCCTAACTTGGTTTTTGTCTGT (for recombination product) |

### Human primers sequence used or QRTPCR

| | Gene Name | Primer Forward (F) and Reverse (R) |
|---|---|---|
| 1 | BMAL1 | F - GCTCAGGAGAACCCAGGTTATC<br>R - GCATCTGCTTCCAAGAGGCTCA |
| 2 | 18S | F - ACCGCGGTTCTATTTTGTTG<br>R – CCCTCTTAATCATGGCCTCA |

**Supplemental Methods:**

***In vitro* synchronization of cells:** Cells were synchronized using 100nM dexamethasone for 2h [1], and released into media containing 5% FBS. 12h after release, cell lysates were collected every 4h, for 48h.

***In vitro* BrdU incorporation:** 25,000 mIMCD3 cells were seeded on glass coverslips and grown in DMEM:F12 media containing 5% FBS and 1% pen/strep. 40-50% confluent cells were exposed to FBS-free media for 16h followed by 0.2% FBS-containing media for 24h. Cells were incubated with BrdU (#10280879001, MilliporeSigma, Burlington, MA) (3 µg/mL) for the last 3 h of incubation. Cells were fixed and immunostained for BrdU (#5292S, Cell Signaling Technology, Danvers, MA) and DAPI. Images were taken using Nikon 80i microscope and quantified using Image J software.

**MTT assay:** To measure the cell viability, 20,000 cells were seeded in 24-well plates. The cells were serum starved overnight followed by treatment with Vehicle or 10 nM Firsocostat (HY-16901, MedChemExpress, NJ, USA) in 0.2 % FBS containing media for 48 hours, and MTT assay was performed as described earlier [2].

**Quantification of cyst:** The kidney sections (5µm) were stained with Hematoxylin and Eosin and images were taken using Nikon 80i upright microscope (Tokyo, Japan), and cyst number, % cystic index and total kidney area were quantified using ImageJ (Fiji, Madison, WI, USA). The number of cysts and its individual area were obtained using the “Measure” function, and the area of all the cysts were added to get the total cystic area. The total kidney area was determined by drawing circumference of the kidney using the “Freehand selections” function which yielded the area using the “Measure” function. Cystic index was calculated by dividing the cyst-containing area by the total kidney area.

**Western blot:**

Mouse kidney tissues were homogenized in SDS Laemmli buffer and loaded onto SDS polyacrylamide agarose electrophoresis gels as described previously [3, 4]. Primary antibodies used were, BMAL1 (14020), pERK1/2 (9101), cyclinD1 (2978 S), S6 (2317), pS6 (4858) p-CREB (9198), CREB(9197) from Cell signaling (Danvers, MA, USA); GAPDH (SC-32233), ERK1/2 (SC-94), p53 (SC-6243), c-Myc (SC-40) from Santa Cruz Biotechnology, Inc. (Dallas, TX, USA). The secondary antibodies for anti-rabbit (P0448) and anti-mouse (P0447) were purchased from Dako (Santa Clara, CA, USA). The ECL reagent was purchased from Amersham (GE Healthcare, Buckinghamshire, UK).

**Immunohistochemistry/immunofluorescence (IHC/IF) staining:**

For IHC, the mouse kidney tissues were fixed in 4% paraformaldehyde and embedded in paraffin as described before [5]. For IHC, the primary antibodies were, BMAL1 (14020) and Ki-67 (#12202) from Cell Signaling Technology, (Danvers, MA). Tissue sections were incubated with streptavidin HRP conjugate secondary antibody (Invitrogen, Carlsbad, CA, USA), followed by DAB (Vector Laboratories, Burlingame, CA, USA). Following counterstaining with Harris Hematoxylin stain, tissues were dehydrated, and mounted using Permount (Fisher Scientific, Waltham, MA, USA).

For IF, DBA (Vector Laboratories, Burlingame, CA), Collagen Type-1a (#203002 from MD Bioproducts, Oakdale, MN), $\alpha$-SMA (#ab5694 from Abcam, Cambridge, MA) primary antibodies were used. Secondary antibodies used were goat anti-Rabbit IgG fluor and goat anti-mouse IgG

Texas red (Invitrogen, Carlsbad, CA, USA). After incubation with secondary antibodies, tissue section were washed and stained with DAPI, and mounted with Flour-G (Invitrogen, Carlsbad, CA, USA). Images were taken using a Nikon 90i upright microscope (Tokyo, Japan).

**TUNEL assay:**

TUNEL assay was performed on mouse kidney tissue sections using an *In Situ* cell death detection kit, Fluorescein (#11684795910 Roche, Mannheim, Germany) as described before [6].

**Quantitative real-time PCR:**

RNA was isolated from mouse kidney tissue or cultured cells using trizol method (Ambion, Austin, TX, USA). High-capacity cDNA reverse transcription kit from Applied Biosystems (# 4368814, Foster City, CA, USA) was used to make cDNA according to the manufacturer's protocol. Quantitative real-time PCR was done using power SYBR Green PCR master mix from Applied Biosystems (Foster City, CA, USA) according to the manufacturer's protocol. The mouse primer list is provided in the Supplemental Table 1. 18S mRNA levels were measured to normalize gene expression. The individual Ct values obtained from RTPCR reaction cycle was used to calculate 2ˆ(–delta delta Ct). The control group (WT or RC/RC) was normalized to 1 and the fold change was represented as a graph for individual genes.

**Triglyceride assay:** The triglyceride levels in the kidney tissue lysate were measured using manufacturer's protocol (ab65336, from Abcam, Cambridge, MA) and normalized by weight of the kidney tissue (mg) for each sample.

**BUN (Blood urea nitrogen):** BUN levels in the plasma were measured using QuantiChrom Urea Assay Kit (BioAssay Systems, Hayward, CA, USA) as per the manufacturer's protocol.
